# Combined statistical-biophysical modeling links ion channel genes to physiology of cortical neuron types

**DOI:** 10.1101/2023.03.02.530774

**Authors:** Yves Bernaerts, Michael Deistler, Pedro J. Gonçalves, Jonas Beck, Marcel Stimberg, Federico Scala, Andreas S. Tolias, Jakob Macke, Dmitry Kobak, Philipp Berens

**Affiliations:** Hertie Institute for AI in Brain Health, University of Tübingen, 72076 Tübingen, Germany; Tübingen AI Center, 72076 Tübingen, Germany; Champalimaud Centre for the Unknown, Champalimaud Foundation, 1400-038, Lisbon, Portugal; Department of Computer Science, University of Tübingen, 72076 Tübingen, Germany; VIB-Neuroelectronics Research Flanders (NERF), Belgium; Department of Computer Science, KU Leuven, 3001, Leuven, Belgium; Department of Electrical Engineering, KU Leuven, 3001, Leuven, Belgium; Sorbonne Université, INSERM, CNRS, Institut de la Vision, 75012 Paris, France; Baylor College of Medicine, Houston, 77030, TX, USA; Department of Ophthalmology, Byers Eye Institute, Stanford University, Stanford, 94303, CA, USA; Department of Empirical Inference, Max Planck Institute for Intelligent Systems, 72076 Tübingen, Germany

## Abstract

Neural cell types have classically been characterized by their anatomy and electrophysiology. More recently, single-cell transcriptomics has enabled an increasingly fine genetically defined taxonomy of cortical cell types, but the link between the gene expression of individual cell types and their physiological and anatomical properties remains poorly understood. Here, we develop a hybrid modeling approach to bridge this gap. Our approach combines statistical and mechanistic models to predict cells’ electrophysiological activity from their gene expression pattern. To this end, we fit biophysical Hodgkin-Huxley-based models for a wide variety of cortical cell types using simulation-based inference, while overcoming the challenge posed by the mismatch between the mathematical model and the data. Using multimodal Patch-seq data, we link the estimated model parameters to gene expression using an interpretable sparse linear regression model. Our approach recovers specific ion channel gene expressions as predictive of biophysical model parameters including ion channel densities, directly implicating their mechanistic role in determining neural firing.

## 1 Introduction

Neural cell types form the basic building blocks of the nervous system^1^. In the neocortex, they form intricate circuits giving rise to perception, cognition, and action^2–4^. Scientists have classically characterized these cell types by their anatomy or electrophysiology, and more recently, using molecular markers^3,5–7^. In the past decade, single-cell transcriptomics has enabled an increasingly fine genetically defined taxonomy of cortical cell types^8–11^, but the link between the gene expression profiles of individual cell types and their physiological and anatomical properties remains poorly understood.

To tackle this question, Patch-seq has been developed to combine electrophysiological recordings, single-cell RNA sequencing (scRNA-seq) and morphological reconstruction in individual neurons^12–15^. This approach has made it possible to directly study the relationship between the gene expression profile of a neural cell type and its physiological and anatomical characteristics. These studies have found that distinct families of neurons (such as *Pvalb* or *Sst* interneurons, or intratelencephalic pyramidal neurons) show distinct physiological and anatomical properties^16,17^. Within these families, cell properties often vary continuously^17^, possibly caused by smooth changes in gene expression.

This wealth of data has led to the development of sophisticated techniques for multimodal data integration and analysis^18–21^, but uncovering the deterministic relationships between e.g. transcriptomic and physiological properties of the neurons has been challenging. For example, sparse reduced-rank regression can reveal patterns of ion channel gene expression statistically predictive of particular expert-defined electrophysiological features^17^, but precludes a potential causal interpretation. Establishing mechanistic links experimentally is challenging as well, as it involves genetic or pharmacological interventions.

Here, we argue that biophysical models of the physiological activity of neurons can help to close this gap as their parameters are explicitly interpretable (Fig. 1, right). We constructed conductance-based models with single compartments^22–24^ for the electrophysiological activity of 955 neurons from adult mouse motor cortex (MOp) spanning various neural types and classes^17^. In contrast to the expert-defined features previously used to relate gene expression and physiological response patterns, the parameters of these models correspond to mechanistically interpretable quantities such as ion channel densities. We then applied sparse reduced-rank regression to predict the fitted conductance-based model parameters from the gene expression patterns in the same set of cells, completing the statistical-biophysical bridge of gene expression to electrophysiological activity with mechanistically interpretable latent quantities (Fig. 1).

**Figure 1.**
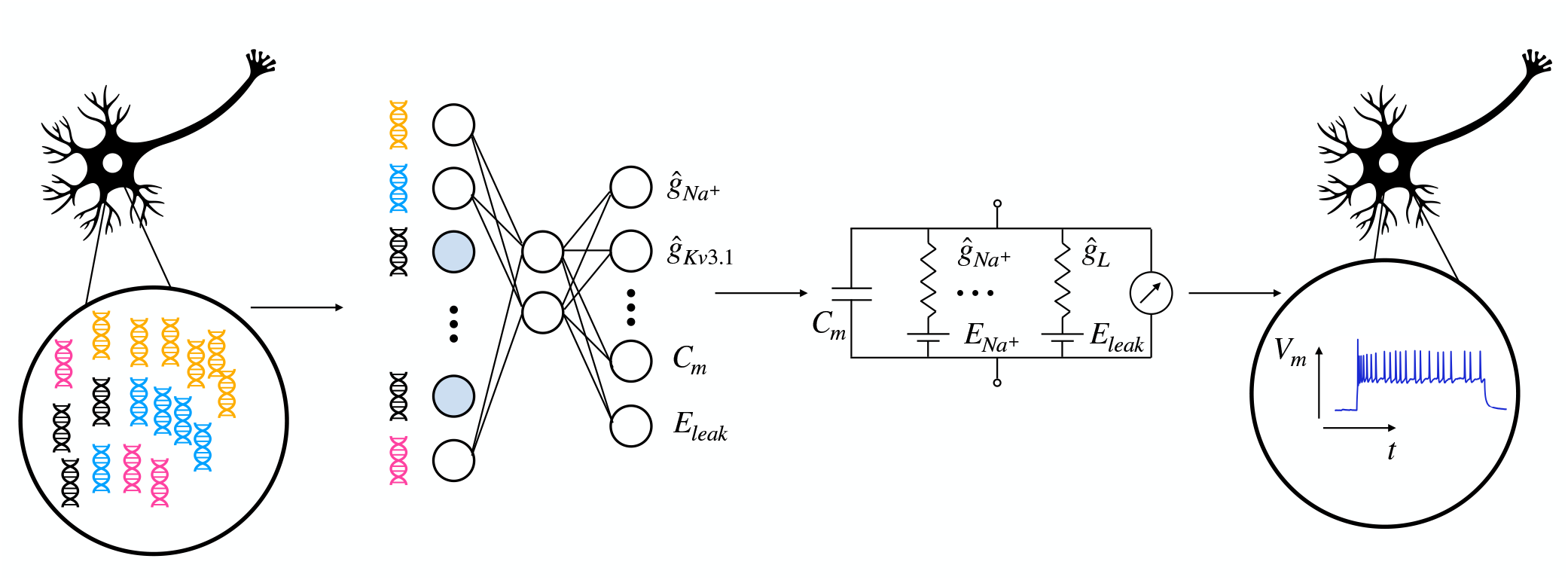
Sketch of the statistical-biophysical hybrid model. Neuronal gene expression levels (left) as well as electrophysiological patterns (right) are obtained experimentally with Patch-seq. The electrophysiology is fitted with a conductance-based biophysical model (middle right). The estimated model parameters are then predicted with sparse reduced-rank regression (middle left), completing the statistical-biophysical bridge from neuronal genotype to phenotype.

In order to find parameters of the mechanistic model that explain observed electrophysiology, we used neural posterior estimation (NPE)^25–27^. This approach can recover the parameters of a mechanistic model based on summary statistics derived from model simulations, providing a posterior distribution over the model parameters. The posterior distribution allows to quantify the uncertainty in our model parameter estimates, in contrast to previous work using genetic algorithms^28,29^. As has been observed in other contexts^30–33^, we found that NPE failed for our biophysical model and our data set due to a small but systematic mismatch between the data and the model, which could not be easily remedied by standard modifications to the model. We developed an algorithm that introduces noise to the summary statistics of model simulations used to train the density network, which allowed it to perform reliable inference despite the model misspecification. We termed it NPE-N, for neural posterior estimation with noise.

With NPE-N, we obtained posteriors over the conductance-based model parameters for all 955 neurons in our diverse data set. We found that parameter samples from the posterior provided for good model fits to the physiological firing patterns of most neurons, but observed higher parameter uncertainty in some families such as *Vip* interneurons. Furthermore, we showed that the relationship between gene expression patterns and the inferred model parameters could be learned using statistical techniques such as sparse reduced-rank regression, allowing to predict the electrophysiology of a cell from its gene expression across cortical neuron types and classes. Our approach recovered specific ion channel genes as predictive of model parameters corresponding to matching ion channel densities, directly implicating them in a mechanistic role for determining specific neuronal firing patterns that differ between cell types.

## 2 Results

### 2.1 Hodgkin-Huxley-based models reproduce Patch-seq electrophysiological recordings

To better understand how the genetic identity of a cortical neuron determines its physiological activity, we studied a previously published data set of neurons from mouse motor cortex, which had been characterized with respect to their gene expression profile and electrophysiological properties using Patch-seq^17^ (Methods 4.1). We focused on a subset of 955 cells which displayed action potentials to injection of a step current of 300*pA* and passed transcriptomic quality control. These neurons had non-zero expression of 7.2 thousand genes on average (ranging between 1.2 and 18.1 thousand genes). The data set included 278 pyramidal neurons and 677 interneurons, consisting of 289 *Pvalb*, 240 *Sst*, 54 *Vip*, and 11 *Sncg* neurons, using the cell families and finer cell type labels assigned by the original authors^17^ based on mapping the gene expression patterns of the neurons to a larger reference atlas^11^.

We hypothesized that we could further clarify the relationship between gene expression patterns and the electrophysiological response properties of the neurons, if we knew the mechanistically interpretable biophysical parameters underlying those. Therefore, we implemented a single-compartment conductance-based model based on a previously established ‘minimal’ Hodgkin-Huxley-based (HH-based) model that captures the electrophysiology in a wide range of neuronal families^23^. We then increased the flexibility of our model with the addition of a small number of ion channels important for modeling pyramidal cells^24^. The parameters of the resulting model included passive parameters such as the capacitance and input resistance as well as active conductances of different ion channels or currents, which determine the physiological responses (Methods 4.2). Our final model included different sodium (*Na*^+^), potassium (*K*^+^), calcium (*Ca*^2+^) and leak currents and had 13 free parameters overall, which needed to be inferred from experimental recordings. We summarized experimental recordings by 23 expert-defined electrophysiological features, including latency to the first spike, action potential count, amplitude and width or the membrane potential mean and variance (for a full list of all 23 features, see Methods 4.3), and computed the same features for each HH-model simulations.

Our HH-based model was able to generate simulations that were close to experimental observations from all major families of neurons, both qualitatively and quantitatively. We defined a uniform prior distribution over the 13 free parameters within biologically plausible ranges, sampled 15 million parameter combinations from it, and ran the biophysical simulation for each of them with the Brian2 toolbox^34^. Out of the 15 million simulations, about 7 million had well-defined values for each of the 23 electrophysiological features. For a given Patch-seq neuron, we picked the parameter combination yielding the simulation lying closest (in terms of Euclidean distance) to the actual neuron in standardized electrophysiological feature space (each feature was Z-scored with the mean and standard deviation of 7 million simulations). This simulation was typically qualitatively similar (Fig. 2) and matched experimental electrophysiological features well (Table 1, last row). However, this strategy required a very large library of precomputed prior simulations and yields only a point estimate, i.e. the best fitting model parameter vector, and no uncertainty information.

**Table 1.** Performance of various NPE training approaches. For the descriptions of training approaches, see Section 4.5. The columns show the percentage of simulation fails (at least one undefined summary statistic) and the Euclidean distance to the experimental values (mean±SD over *n* = 955 MOp neurons) using MAP parameters and using 10 randomly drawn samples from the NPE posterior. In the last row, we take the prior simulation closest to the experimental observation and sample 10 parameter combinations from the prior. Bold values show best rows in each column (excluding the last row).

| Training | MAP |  | Posterior |  |
| --- | --- | --- | --- | --- |
|  | Fails (%) | Eucl. distance | Fails (%) | Eucl. distance |
| Standard NPE | 12.77 | $6.24 \pm 3.24$ | 17.69 | $6.63 \pm 3.24$ |
| Best Euclidean | 14.14 | $5.11 \pm 3.20$ | 20.86 | $5.70 \pm 3.36$ |
| Add 0.001 amplitude noise to ephys | 7.33 | $5.36 \pm 3.10$ | 12.58 | $5.43 \pm 3.17$ |
| Add 0.01 amplitude noise to ephys | 5.86 | $4.81 \pm 2.96$ | 8.54 | $5.39 \pm 3.11$ |
| Add 0.05 amplitude noise to ephys | 4.61 | $4.71 \pm 2.98$ | 9.01 | $5.34 \pm 3.06$ |
| Add 0.1 amplitude noise to ephys (NPE-N) | 2.51 | $4.35 \pm 2.86$ | **6.22** | **5.11 ± 3.03** |
| Add 1 amplitude noise to ephys | **1.36** | **3.81 ± 2.11** | 7.27 | $5.19 \pm 2.56$ |
| Add 0.05 amplitude noise to ephys and model parameters | 3.66 | $5.42 \pm 2.91$ | 10.28 | $6.31 \pm 2.86$ |
| Data augmentation | 3.35 | $4.47 \pm 2.98$ | 6.98 | $4.91 \pm 3.12$ |
| Prior | 0.00 | $2.63 \pm 0.81$ | 52.29 | $11.79 \pm 3.31$ |

**Figure 2.**
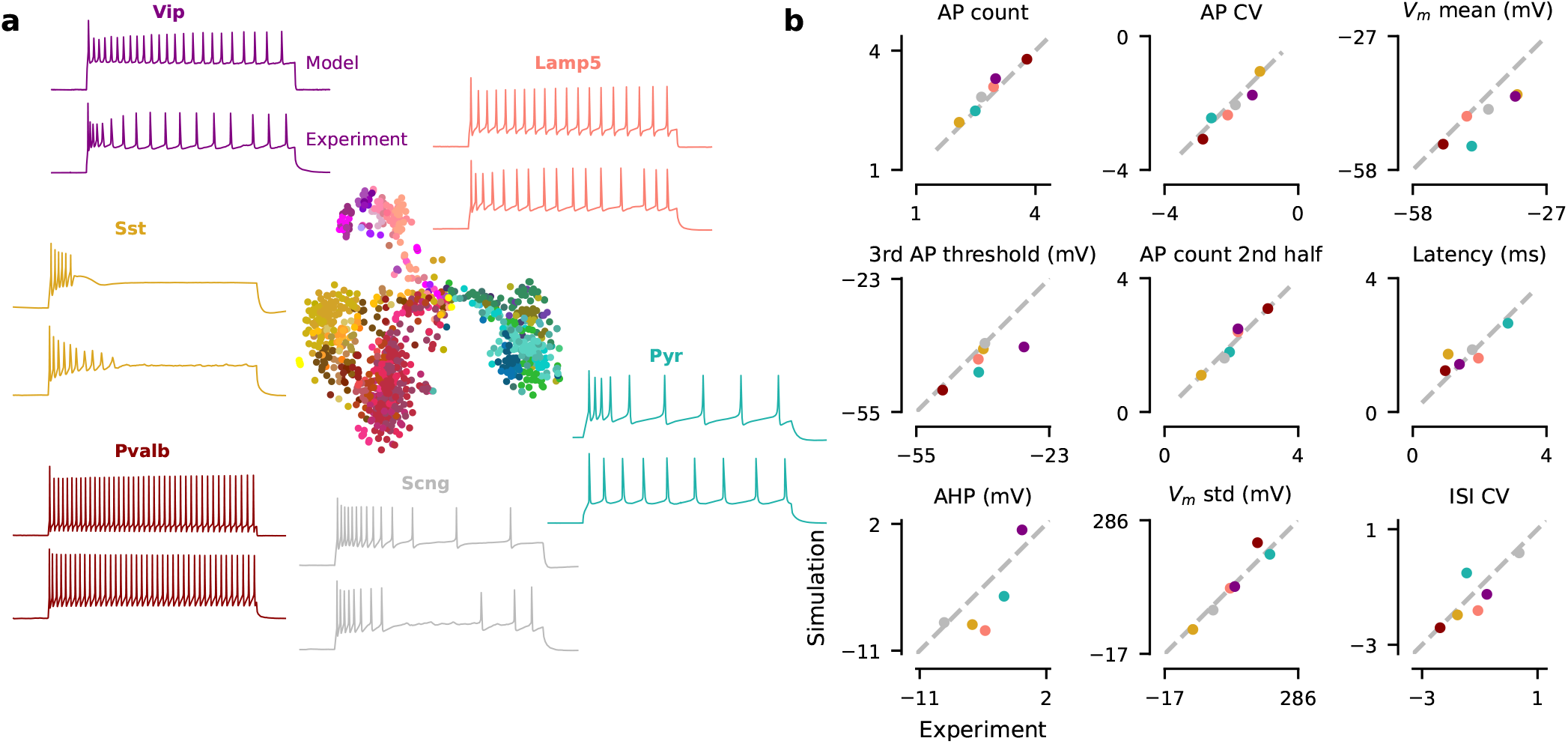
Exemplary experimental observations and their closest simulations from the prior. **a** Middle: t-SNE embedding of *n* = 955 MOp cells based on their transcriptome. Surround: one example neuron for each of the six transcriptomic families. For each neuron, we show the experimental observation (below) and the biophysical model simulation from the prior with the smallest Euclidean distance in standardized electrophysiological feature space (above). **b** Comparison of nine electrophysiological feature values between experimental observations and best prior simulations shown in (a).

### 2.2 Neural posterior estimation with noise

Therefore, we used neural posterior estimation (NPE)^25,26,35^, an algorithm for simulation-based inference^36^, which learns an approximate posterior distribution *q (θ* | **x**) over the parameter vector *θ* given a vector of features **x** computed from the experimental recording (Methods 4.4). The posterior distribution is parameterized as a sufficiently flexible neural density estimator. In contrast to previous methods^28,29^, this approach allowed us to quantify the uncertainty in the parameters after seeing the data. We trained the neural density estimator on our synthetic data set comprising 7 million simulations from our HH-based model to infer the 13 free parameters given the 23 electrophysiological features. After training, the neural network could be evaluated on features of *any* experimental recording and returned the corresponding posterior distribution without further simulations or training, providing a model of this relationship.

However, this procedure did not work well for the neurons in our data set. In many cases, samples from the posterior or the maximum-a-posteriori (MAP) parameters did not produce simulations that came close to the experimental data (Fig. 3). Moreover, 18% of posterior-sampled parameters produced simulations with at least one undefined summary feature, for example undefined latency due to complete lack of action potentials (Table 1). We investigated the reason for this failure and found that the poor performance of the inference framework was due to a systematic mismatch between the electrophysiological recordings and simulations from the model (Fig. 4a), a phenomenon recently observed also in other settings^30–33^. For a given simulation, only few other simulations were very close to it in the feature space, but even simulations lying further away still produced qualitatively very similar outcomes (Fig. 4b, orange). In contrast, for a given experimental trace, the distance to the closest simulation was much larger, even when qualitatively the fit looked reasonable (Fig. 4b, blue).

**Figure 3.**
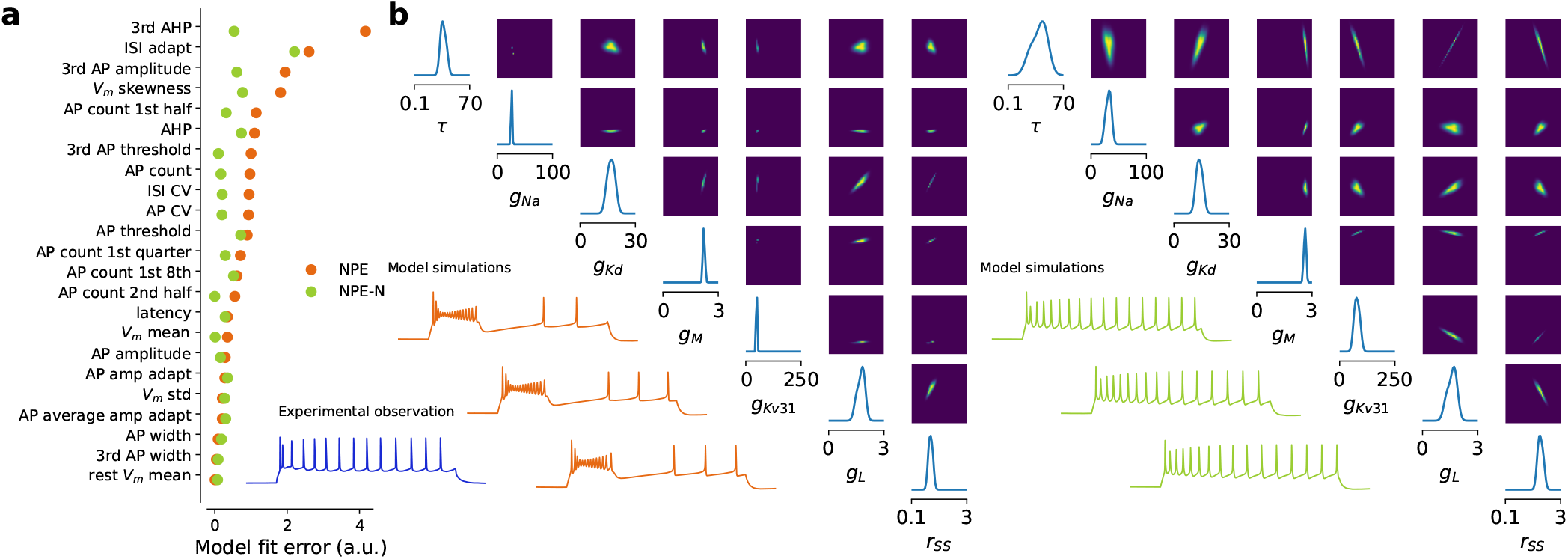
Neural Posterior Estimation vs Neural Posterior Estimation with Noise. **a** The MAP parameter set simulation derived with NPE-N is closer to the experimental reference (in blue below, *L4/5 IT_1* pyramidal cell) than derived with NPE. Residual distance of the MAP parameter set simulation to the experimental observation (model fit error) shown for each electrophysiological feature (0 corresponds to a perfect fit). **b** 1- and 2-dimensional marginals together with 3 simulations generated from parameter combinations with highest probability under the posterior (out of 10 000 samples); NPE (left) vs NPE-N (right) setting. 7 out of 13 model parameters have been selected for illustration.

**Figure 4.**
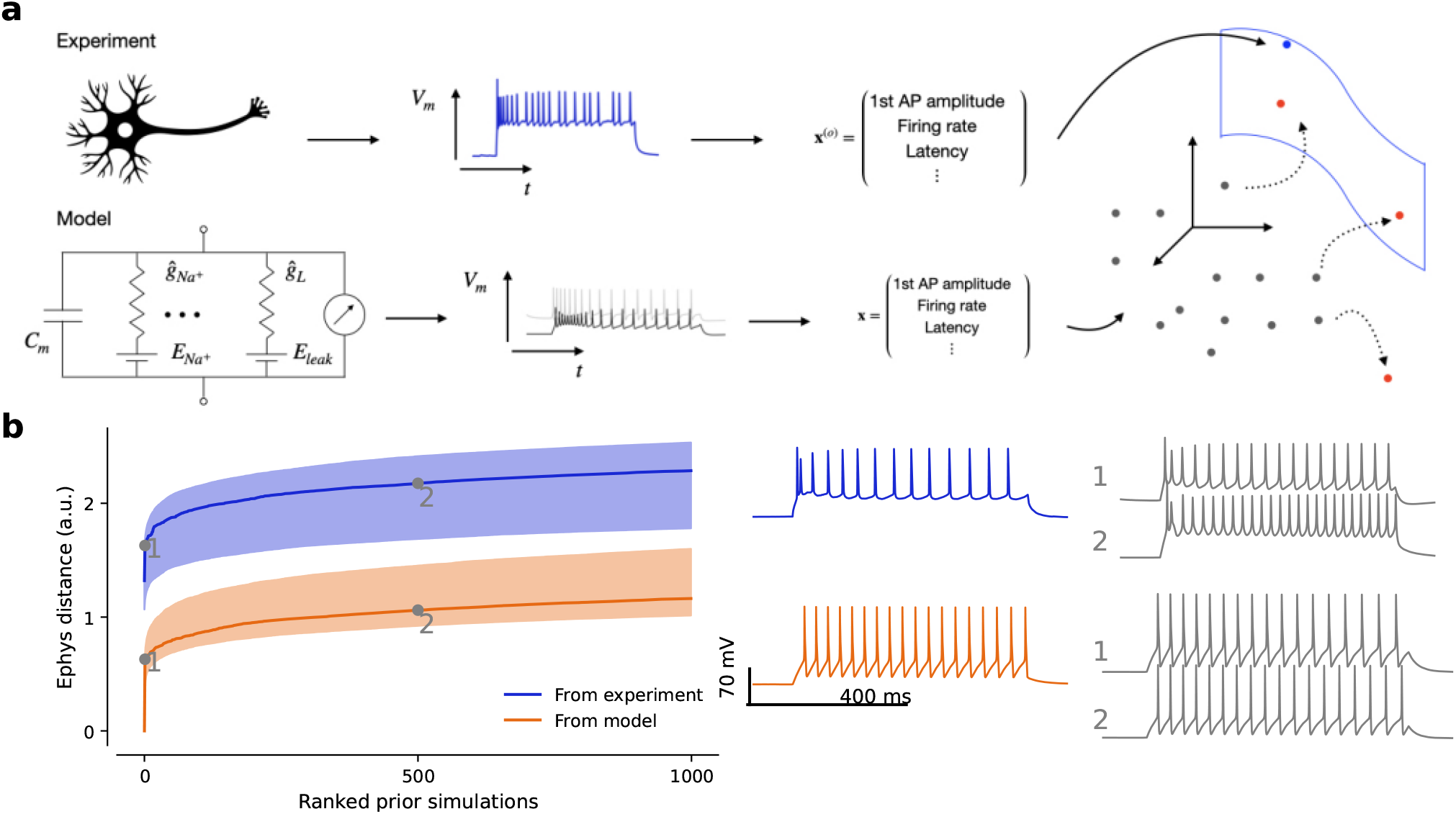
Neural posterior estimation of conductance-based model parameters in the presence of model misspecification. **a** Sketch illustrating model misspecification: in electrophysiological feature space, not enough simulations cover the space of experimental observations. NPE-N introduces isotropic noise to the summary statistics of simulations (dotted arrows). **b** Simulations are further away from experimental observations (blue) than from other simulations (orange). Qualitatively, simulations increasingly further away from an experimental observation look more dissimilar than from another simulation. Numbers 1 and 2 refer to the 2nd and 501st closest simulations respectively.

Our conclusion is that the experimental observations occupied a region of the electrophysiological feature space which was systematically shifted from the region occupied by the model simulations (Fig. 4a), despite our best efforts to create a HH-based model that has sufficient flexibility to capture the response diversity of the cortical neurons in our data set (Fig. 2). For instance, we introduced the *r*_*SS*_ parameter to the model (Methods 4.2), that can scale the speed with which ion channels reach open steady states, in order to alleviate model-data mismatches observed in the action potential width, which were especially large in pyramidal cells and difficult for the model to capture without it. Although this approach substantially reduced overall model mismatch, it did not remove it entirely (Fig. S1). As a consequence, many simulations produced with randomly drawn posterior parameters had either one or more undefined electrophysiological feature or found themselves at a large distance to their experimental reference (Fig. 3a; Table 1).

We found that modifying the neural posterior estimation algorithm improved the performance of the inference procedure. To allow the density estimator to generalize better to experimental observations at test time, we smoothed the feature space by adding a small amount of independent Gaussian noise to the electrophysiological features of selected simulations close to the experimental observations and used those to train the neural density estimator (Fig. 4a and Methods 4.5). This procedure yielded posterior simulations which came much closer to their experimental reference both qualitatively and quantitatively (Fig. 3). The Euclidean distance in electrophysiological feature space between the experimental recording and the simulation with the MAP biophysical parameters ∥**x**_MAP_ −**x**∥ was 4.35±2.86 when using noise vs. 6.24±3.24 when using standard NPE (mean±SD across *n* = 955 cells; Table 1). We experimented with several strategies of adding noise of different magnitude and chose a compromise between simulations from the posterior being close to the measured data and a low fraction of simulations that result in undefined features (Methods 4.5). We called the resulting procedure NPE-N and used it to obtain posterior distributions over parameters for all 955 neurons in our data set. NPE-N outperformed NPE both qualitatively and quantitatively across various cell types as showcased for six additional cells representative of different cell types in Figs. S2–S7.

To gain further insights into the inference procedure, we asked which of the 23 features were most important to constrain the posterior. To this end, we used an algorithm that efficiently compares a posterior constrained by the full set of features to one constrained by a growing subset of features^37^ and studied a subset of 50 neurons (Methods 4.6). We ran this algorithm five times for each of these neurons, and counted how often a feature was selected as one of the five most important features (Fig. S8a). We then compiled the results of this selection procedure across all 50 neurons and found that on average, the mean resting membrane potential was by far the single most important feature, followed by the mean potential during current stimulus, action potential amplitude, the action potential threshold, and the variance of the membrane potential (Fig. S8b).

### 2.3 Transcriptomic, electrophysiological and HH-based model parameter variability

We next returned to our original question and studied how the transcriptomic identity of the neurons in our data set was related to their electrophysiological properties and the MAP parameters of the best-fitting HH-based model. To this end, we used a two-dimensional *t*-SNE visualization of the gene expression data of all 955 MOp neurons (Fig. 5a). We found that the embedding separated the major neural families, including interneurons and pyramidal neurons, well (Fig. 5a). We confirmed the identity of these families by overlaying the expression strength of various marker genes such as *Pvalb, Sst, Vip* and *Lamp5* (Fig. 5b). The NPE-N posteriors for neurons from some families were less constrained than those of others, indicated by higher posterior entropy (Fig. 5c). Specifically, this affected *Vip* neurons, which were relatively sparsely sampled in the data set. In contrast, *Pvalb* neurons showed the lowest uncertainty indicating that their posteriors were best constrained using the available features. One reason for this may be that *Pvalb* neurons fired more stereotypically, whereas *Vip* neurons showed greater variability in their firing patterns^17,38^, that may require greater flexibility in the model to reproduce.

**Figure 5.**
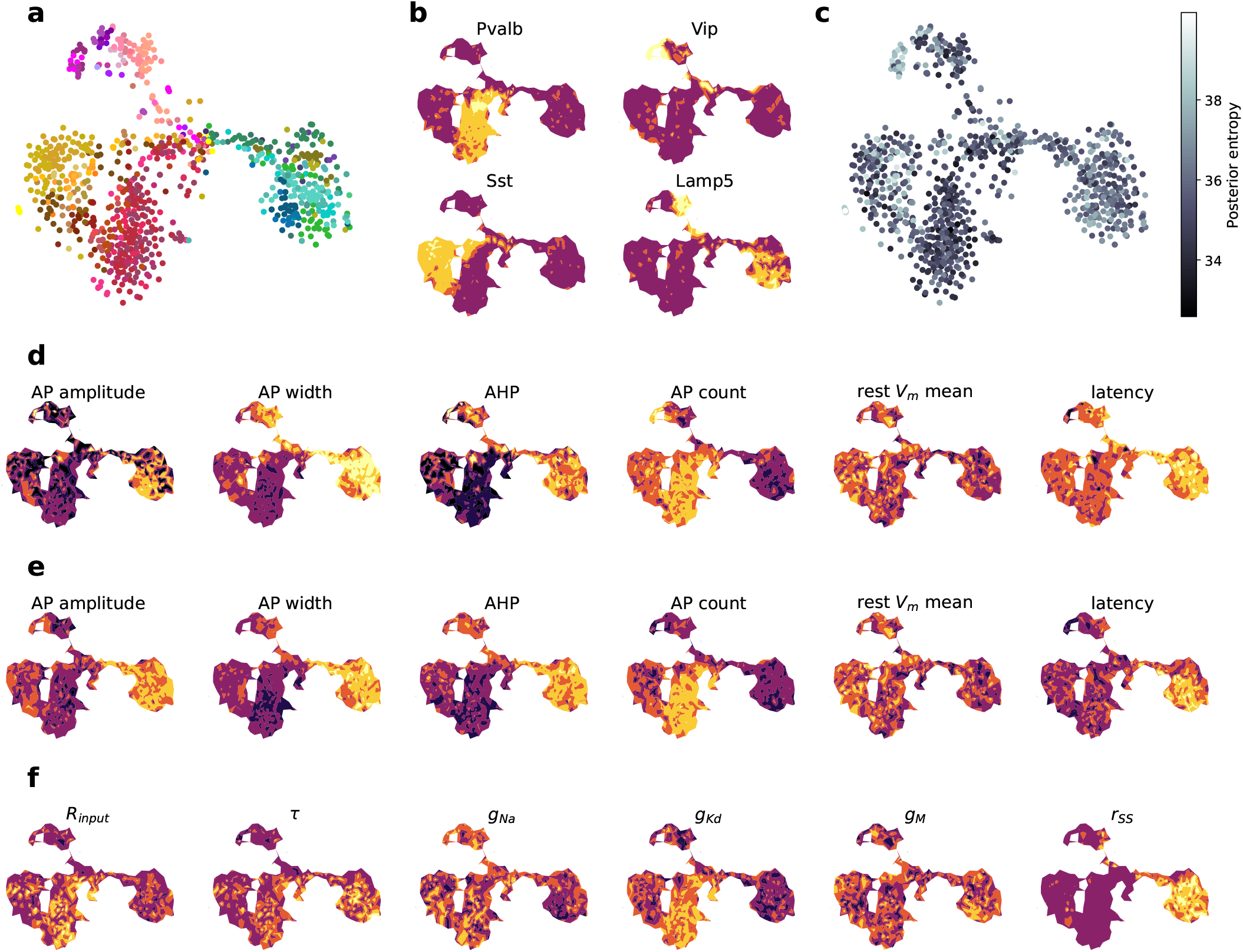
Two-dimensional embedding reveals difference in HH-based parameters between neural families. **a** T-SNE embedding of *n* = 955 MOp neurons based on transcriptomic data. Colors like in Fig. 2a. Cells in the middle of the embedding had lower quality transcriptomic data and therefore grouped together. **b** Marker genes expression levels overlayed and interpolated on embedding confirm known families (dark purple: low expression, yellow: high expression). **c** Uncertainty of MAP parameters for each cell overlayed on the embedding. The uncertainty was calculated as the posterior entropy 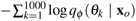, where we sampled *θ*_*k*_ *q*_*ϕ*_ (*θ* **x**_*o*_). **d** Selection of summary statistics derived from simulations corresponding to MAP estimates, overlayed on the embedding. **e** Selection of summary statistics describing observed electrophysiology, overlayed on the embedding. **f** Selection of MAP parameters, overlayed on the embedding.

We overlaid the individual electrophysiological features on this two-dimensional embedding, both for the simulated MAP traces and experimentally measured data (Fig. 5d and e). We found that, as expected, these features varied strongly between neural families, and that the features extracted from simulated traces matched well to the features from measured traces. For example, pyramidal neurons showed higher action potential amplitude and width as well as lower spike rates, in line with the observations. For some features, such as the latency of the response, the match was less perfect, as the simulated traces of interneuron families had overall higher latency than those experimentally measured.

Next, we studied how the parameters of the HH-based model varied across this transcriptomically defined embedding (Fig. 5f). This visualization allowed us to reason about the relationship between biophysical parameters and the resulting electrophysiological properties in some of the genetically defined families. For example, the conductivity of the delayed rectifier potassium current 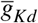^39^ was estimated to be high for *Pvalb* interneurons (Figure 5f), suggesting that these currents were important for quickly repolarizing the membrane potential *V*_*m*_(*t*) during AP generation in order to obtain the small *AP widths* and high *AP count* of fast-spiking *Pvalb* cells (Figure 5d–f). Likewise, the membrane time constant *τ* and the scaling parameter *r*_*SS*_ were important to fit the large action potential widths and high latency observed for pyramidal neurons (Figure 5d–f).

### 2.4 Closing the gap: from genes to electrophysiology

Given these results, we were now in a position to quantitatively study the relationship between the transcriptomic identity of a neuron and its biophysical parameters. To this end, we trained a sparse linear reduced-rank regression model (sRRR)^18^ and a sparse nonlinear bottleneck neural network (sBNN)^19^ to predict the biophysical parameters (*d* = 13) from the gene expression data (Fig. 6a). To ease the interpretability, we focused on ion channel and known marker genes only (*d* = 427) and trained linear and nonlinear models with a two-dimensional latent space. We found that model parameters could be predicted with reasonable accuracy (sRRR: *R*^2^ = 0.17±0.03, mean±SD across cross-validation folds, for a model selecting approximately 25 genes) and that the nonlinear model performed just as well as the linear one (sBNN: *R*^2^ = 0.17±0.03), so we analyzed only the linear model further (Fig. 6b). Over the entire data set, this model predicted some parameters such as the conductance of the fast non-inactivating and delayed rectifying potassium channel (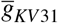and 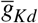) or the membrane capacitance *C* particularly well (Fig. 6c). Other model parameters were less well predicted, such as the leak potential *E*_*leak*_ or the muscarinic potassium channel conductance 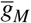. Interestingly, the *r*_*SS*_ parameter, which we introduced as the first step towards alleviating model mismatch issues, was predicted the best.

**Figure 6.**
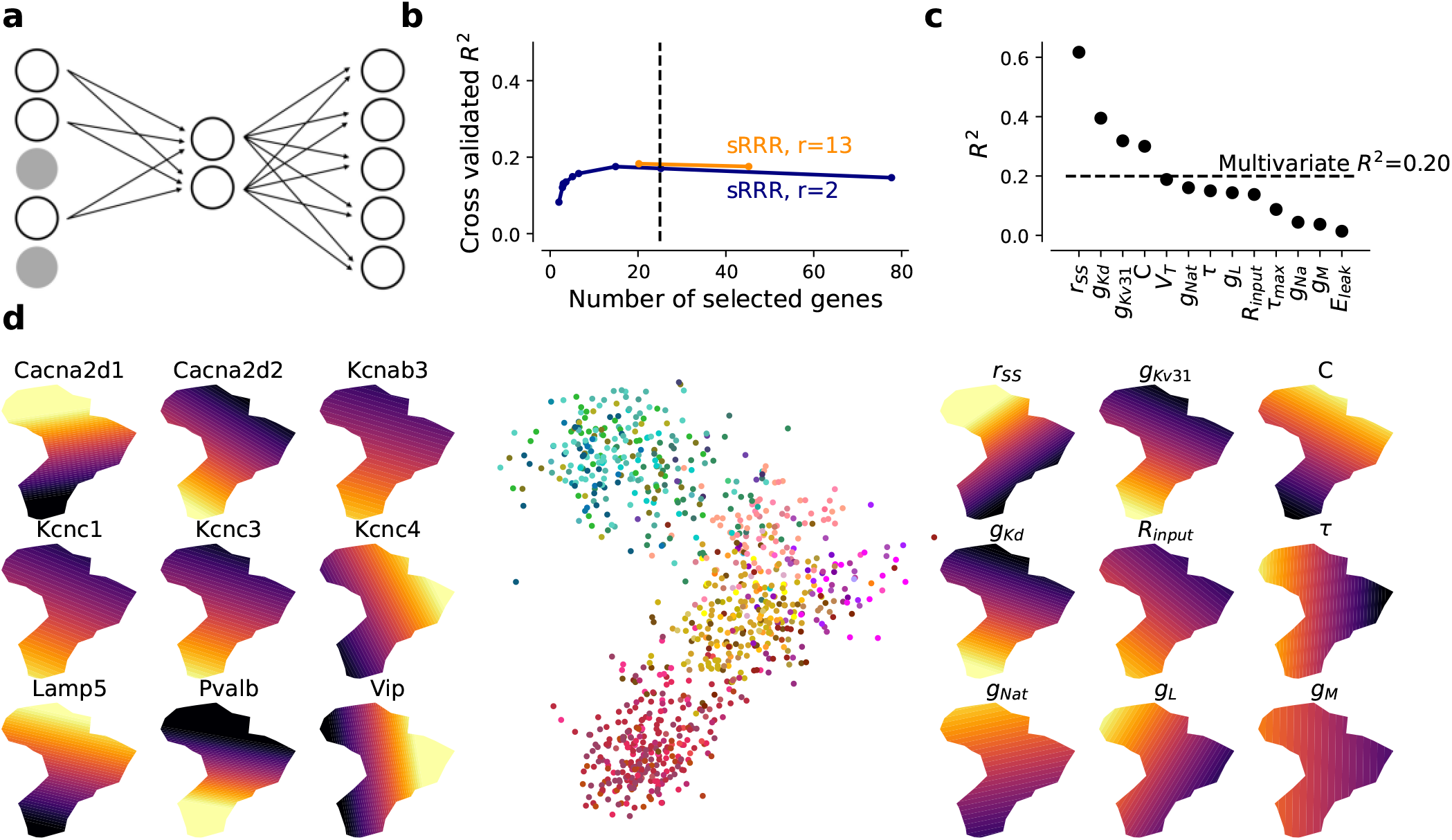
Prediction of MAP parameter estimates from gene expression with sparse reduced-rank regression. **a** sRRR schematic. A linear combination of selected genes is used to predict fitted HH-based model parameters. **b** Cross-validation performance for rank-2 and full-rank sRRR models with elastic net penalty. The dashed vertical line shows the performance with 25 genes. **c** Rank-2 sRRR model predictive performance for each model parameter, using the entire data set. **d** Middle: rank-2 sRRR model latent space visualization. All 955 MOp neurons are shown. Left: Selected ion channel and marker gene overlays. Right: Predicted model parameter overlays.

We visualized the latent space of the sRRR model to better understand the relationship between ion channel and marker genes and the HH-based model parameters (Fig. 6d). This embedding is conceptually similar to the *t*-SNE visualization of the entire gene space with overlayed model parameters and electrophysiological properties (Fig. 5), except that here we focus on genes that predict HH-based model parameters. The 2D latent space of the sRRR model showed two principal directions of variation, where one separated pyramidal cells from interneurons and the other distinguished different interneuron families. In addition, we found that the sRRR model identified mechanistically plausible relationships: for example, the potassium channel conductances 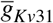 and 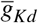 were both high in *Pvalb* neurons placed in the lower left corner, predicted by the expression of various potassium channel genes like *Kcnc1*, that constitutes a subunit of the Kv3.1 voltage-gated potassium channel, and *Kcnab3* respectively. Likewise, the calcium channel conductance 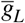was predicted by high expression of *Cacna2d1* that directly encodes for the alpha-2 and delta subunits in the L-Type calcium channel. Our sRRR model selected *Cacna2d2* as well, which is a paralog gene with opposite expression to *Cacna2d1* (Fig. 6d, left). In addition, classical marker genes like *Vip* acted as surrogate cell family markers and contributed to the prediction.

This approach can be used to predict HH-based models for neurons for which we only measured gene expression but not electrophysiology, especially on the family level. The model predictions captured essential variation in the model parameters on the family level, although less so on the cell-type level (Fig. 7a, b). To quantify this, we measured the Euclidean distance (normalized by variance) between the matrix with average NPE-fitted MAP parameter values and sRRR-predicted parameter values (Fig. 7a and b, respectively). On the family level (Fig. 7a and b, left panels), this distance was substantially smaller than on the cell-type level (Fig. 7a and b, right panels) — 18.02 (family level, left panels) vs 50.15, 41.94, 103.21, 118.29, 28.70 and 126.02 (for *Lamp5, Sncg, Vip, Sst, Pvalb* interneurons and pyramidal cells respectively, right panels) — indicating that the variation in gene expression levels can be used to predict electrophysiology accurately on a family level, but less so on the cell-type level. We made a qualitative comparison by simulating the HH-based model first with the average MAP estimate across families and then with the average family-based sRRR model prediction (i.e. using only gene expression levels). Except for *Scng* interneurons for which we had only few (*n* = 11) cells available, sRRR-based predictions matched the electrophysiological feature values of MAP-based model predictions and generated simulations almost indistinguishable by eye (Fig. 8).

**Figure 7.**
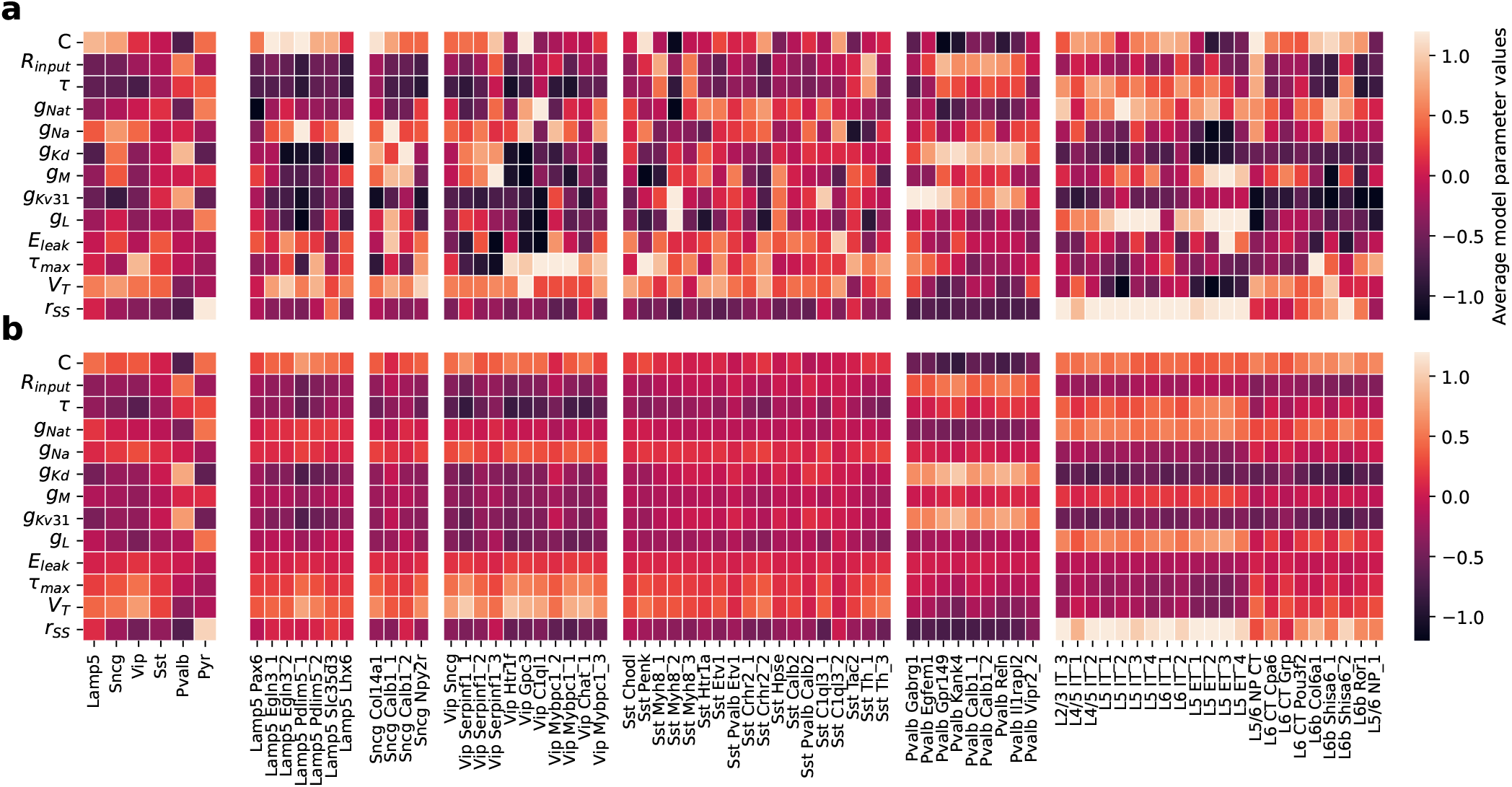
MAP parameter estimates and sRRR predictions for each family and cell type. **a** MAP parameter estimates averaged over cells belonging to a family and belonging to a transcriptomic cell type (left and right respectively). We Z-scored all values by subtracting and scaling with the mean and standard deviation of *n* = 955 MAP estimates, respectively. **b** Analogous to a, but with rank-2 sRRR predicted model parameter values.

**Figure 8.**
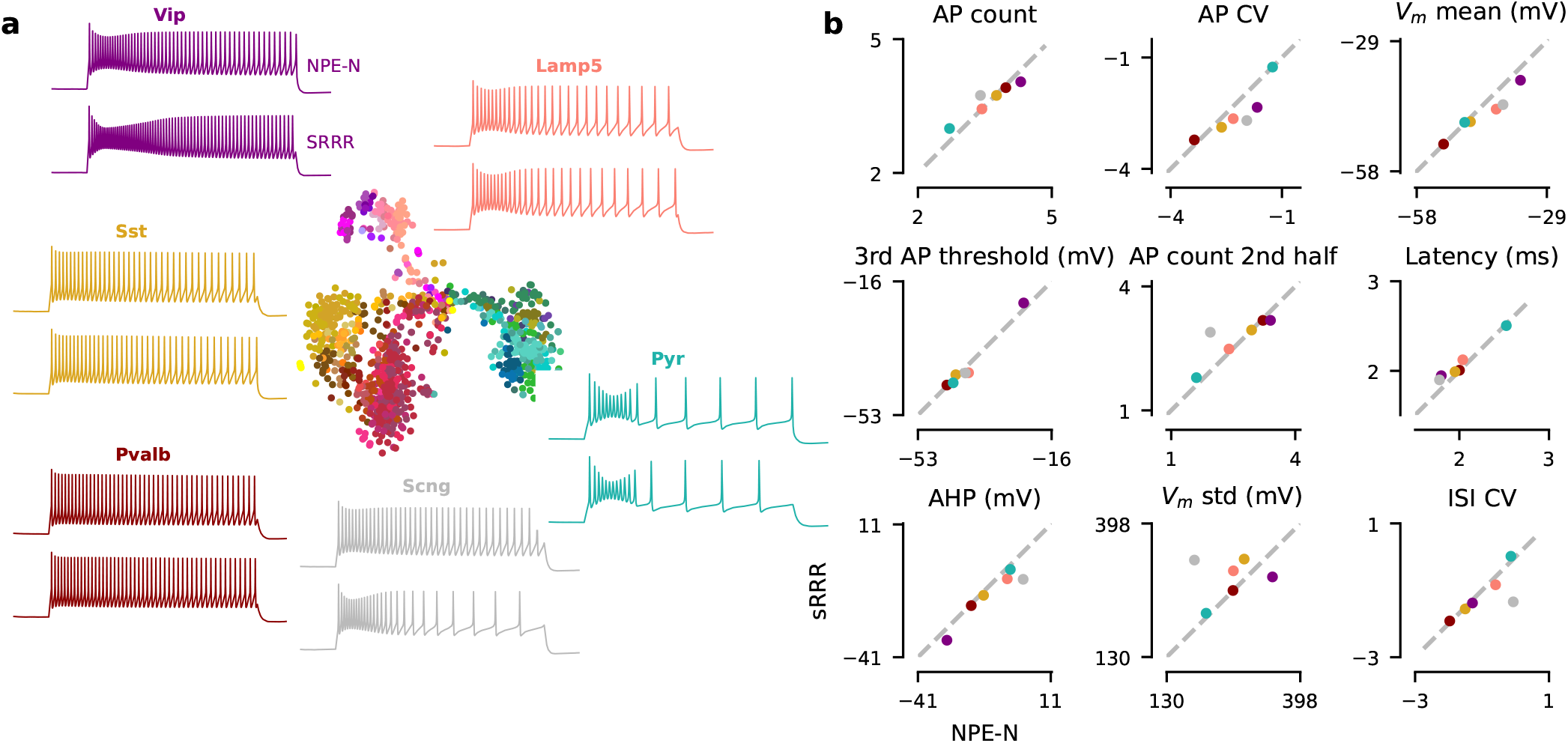
Family representation of MAP estimates together with sRRR predictions. **a** Analogous to Fig. 2a except that the simulation on top is derived from the family-average MAP estimate calculated as in Fig. 7a, left. Simulation on the bottom is derived from the family-average sRRR prediction calculated as in Fig. 7b, left. **b** Comparison of 9 electrophysiological feature values derived with the MAP estimate versus sRRR-based estimate.

## 3 Discussion

In this study, we directly linked the gene expression profile of a set of cortical neurons to HH-based model parameters fitted to their electrophysiological signatures. We believe this is a major step towards closing the causality gap between gene expression and neural physiology: In previous work, we and others have simply correlated gene expression and electrophysiology, e.g. predicting electrophysiological properties from transcriptomic data^17–21^, but uncovering the causal relationships between these quantities has been challenging.

The mechanistic HH-based model we used here spells out our causal understanding of how electrophysiological properties arise from ion channel densities and other passive properties of a neuron, leaving the link between these quantities and the expression of certain genes to be explained by a statistical model. In our approach, we used a linear reduced-rank regression model with groupwise sparsity constraint on the genes, selecting genes in or out of the model based on the available data. Given the present data, we found that the linear model with a two-dimensional intermediate layer performed as well as a comparable nonlinear model. Partially, this may be due to the noise in gene expression and the comparably small data set, but it is also possible that our explicit mechanistic model for the generation of electrophysiological activity explained away some of the nonlinear relationship between gene expression and electrophysiological features. Larger data sets will likely also help the linear model to resolve better observed differences in model parameters between fine cell types, which are currently captured only to a certain extent (Fig. 7).

The *r*_*SS*_ parameter was introduced to improve the used HH-model for the wider *AP widths* observed in pyramidal cells and significantly alleviated some of the model mismatch that we originally observed (Fig. S1). This parameter adapts the rate with which *Na*^+^ and *K*^+^ ion channel gates reach their respective *steady state* (SS) in the model (Table 2), effectively changing the dynamics of in-and outflow of these ions and therefore the *AP width*. Yet, it is possible that additional biophysical factors could account for this, such as more complicated morphology, which may be necessary to capture pyramidal neuron physiology.

**Table 2.**
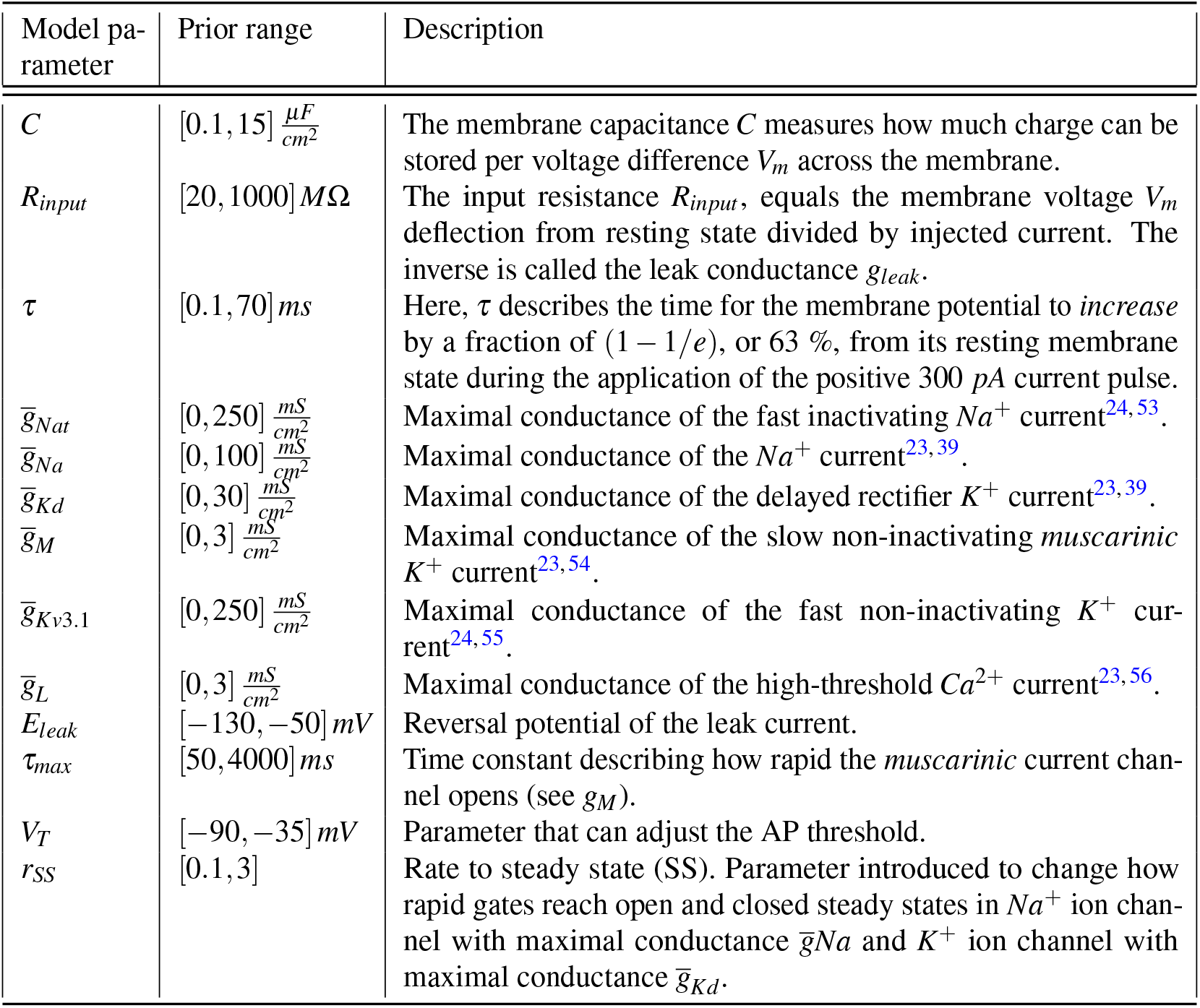
Description of the 13 HH-based model parameters.

Previous work has also attempted to infer biophysical parameters in HH-based models and link the inferred values to neural cell classes and their gene expression^23,24,28,29^. Unlike our work, most of these studies did not directly link parameters in HH-based models to the expression of a large set of genes, but rather studied parameter differences between genetically defined cell classes^23,24,28^. One recent study examined the relationship between HH-based model parameters and individual genes^29^, but did not provide a systematic statistical model for this link. Also, none of these studies used uncertainty-aware parameter inference techniques. Incorporating uncertainty allowed us to highlight cells, cell types or classes for which the inference procedure returned results which were not as well constrained by the data.

Nevertheless, many trends in model parameters across different cell classes qualitatively matched previous observations. For instance, we found that different values are needed for the potassium conductance 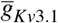to model *Vip, Sst* and *Pvalb* neurons and that the expression of *Kcnc1* varies accordingly (Fig. 6d). In a similar vein, Nandi et al. report different values for 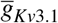 and show that *Kcnc1* is differentially expressed between these classes^29^. In their work, this example is hand-selected for analysis, while in our analysis the evidence emerges from the sRRR model. While not all correspondences with other studies are as close as this, HH-based models show redundancy in their parametrizations^25,40,41^, such that different model parameter combinations can result in similar responses, potentially explaining observed deviations.

Our biological question to identify a quantitative link between gene expression and electrophysiological activity patterns led us to devise a methodological innovation to overcome the challenges posed by systematic mismatch between our simulations and the experimental data, which caused the out-of-the-box simulator-based methods to fail. We achieved this by adding noise to the summary statistics derived from the simulations, effectively smoothing the feature space. In parallel work to this paper, this phenomenon has recently received more widespread attention^30–33^: for instance, robust neural posterior estimation^30^ takes the opposite approach to our strategy and denoises the measured data towards the model using Monte-Carlo sampling. On the other hand, robust synthetic likelihood approaches^42^ that estimate likelihoods rather than posteriors, work similarly to our approach. Which strategy works best for which models and circumstances remains to be evaluated, but these strategies will allow to apply simulation-based inference techniques in cases where models provide relatively coarse, but useful approximations of the true phenomena. Alternatively, one could make the model more realistic. In our case, some of the model mismatch is likely also caused by the use of single-compartment models in contrast to other studies, which used HH-based models with two or more compartments^24,28,29^, however, such complex models are currently difficult to use with simulation-based inference.

We could not predict the electrophysiology of each cell type, based on their average gene expression levels as well as we could for each family of neurons. While this predictive power varied between families, some important patterns were clearly missed by the model. Likely, this is due to transcriptomic data being noisy on the single-cell level^43^. Further, the size of the dataset was quite limited given the number of different cell types, with less than 14 neurons per cell type on average to learn from. Furthermore, covariablity between transcriptomic and physiological parameters may be more continuously changing within one family^17^, making cell type level predictions even harder.

Mechanistic models that make biophysical processes explicit are ultimately desirable all the way from gene expression to electrophysiology, as such models form the highest level of causal understanding^44^. To further close this causality gap would require an explicit mechanistic model for the translation of mRNA into proteins such as ion channels — a relationship, which is all but simple^45–47^. Mechanistic models for this process have been suggested in the literature on a variety of scales^48,49^, but it is an open question how such models could be integrated in the inference procedure given the temporal and spatial processes involved in mRNA translation^48–50^. While directly measuring translation dynamics in live cells has become possible using live cell imaging approaches^51,52^, it remains an extremely challenging task, especially in multicellular systems and given the diversity of cortical neurons. Therefore, the combination of machine learning-based models with explicit mechanistic models may provide a viable path forward to improve our understanding of neuronal diversity even if we do not have full causal empirical knowledge of the entire chain of events, aiding the inference of important intermediate quantities.

## 4 Methods

### 4.1 Data set

We reanalyzed a published data set consisting of *n* = 1328 adult mouse motor cortex (MOp) neurons^17^, which had been characterized transcriptomically and electrophysiologically using Patch-seq. We downloaded the read count data from https://github.com/berenslab/mini-atlas and the electrophysiological traces from https://dandiarchive.org/dandiset/000008. The authors used Smart-seq2 to obtain single-cell transcriptomes for these neurons. Out of *n* = 1328 cells, *n* = 1213 cells passed transcriptomic quality control and were assigned a transcriptomic cell type using the 1000 genes that were most variable across this subset of cells. For electrophysiological characterization, the authors injected negative to positive constant currents for 600 ms time windows starting at −200 pA with steps of 20 pA to positive currents beyond 400 pA or until the cell died. Electrophysiological experiments were performed at a temperature of 25 °*C*. For further experimental details, see^17^. Finally, out of *n* = 1328 cells, we analyzed *n* = 955 cells that had well-defined summary statistics in their membrane voltage response to current injection of 300 pA.

### 4.2 Hodgkin-Huxley-based model

We used a single-compartment HH-based model^23^ that was designed to reproduce electrophysiological behavior of a wide variety of neurons across species with a minimal set of ion channels. To account for the variability across excitatory and inhibitory cortical neurons, we added additional ion channels^24^ and introduced *r*_*SS*_, a parameter influencing how rapid gates reach open and closed steady states in some sodium and and potassium currents. Without these modifications we could not fit wider *AP widths* observed in pyramidal cells.

The HH-based model solves the following ODE *V*_*m*_(*t*) = *f* (*V*_*m*_(*t*), *θ*) for *V*_*m*_(*t*), the membrane voltage as a function of time:

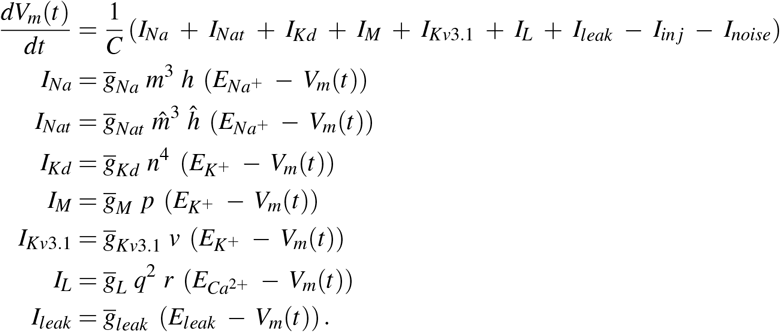

Here, 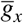and *E*_*x*_ denote the maximum channel conductance and reversal potential of membrane ion channel *x* respectively. *C* is the membrane capacitance and *I*_*in j*_ = 300 *pA* denotes the magnitude of experimental current injected between current stimulation onset at 100 *ms* and stimulation offset 700 *ms*. In order to model small membrane voltage fluctuations observed experimentally, we further introduced Gaussian current noise *I*_*noise*_ ∼ 𝒩 (10 *pA*, 1 *pA*) at every time point.

Furthermore, ion channel activation and inactivation gates follow dynamics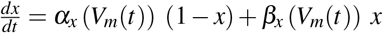, where *x* ∈ {*m, h*, 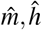, *n, p, v, q, r*}. Opening *α*_*x*_ and closing *β*_*x*_ rate constants depend on the membrane voltage *V*_*m*_(*t*) as previously described^23,24^. In order to account for the 25 °*C* temperature at which Patch-seq experiments were performed, we used a temperature coefficient *Q*_10_ = 2.3 to scale the kinetics with which gates in ion channels open and close. Parameter *r*_*SS*_ further scales the rates with which sodium and potassium currents with maximal conductances 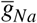and 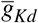reach steady states.

In total, 13 parameters in the model can be tuned in order to fit observed electrophysiology of *n* = 955 MOp cells (Table 2 for details). We implemented the model with the `Brian2` toolbox^34^ in `Python`, which can efficiently transpile and simulate models in *C++*.

In order to understand the significance of adding the *r*_*SS*_ in reducing model misspecification on a “model implementation” level rather than training the density estimator, we compared to the same HH-based model but with scaling parameter *r*_*SS*_ = 1 (Fig. S1).

### 4.3 Electrophysiological features

We automatically extract 23 electrophysiological summary features from the experimental or simulated voltage traces *V* (*t*) using a Python library from the Allen Software Development Kit (SDK) (https://github.com/AllenInstitute/AllenSDK) with modifications to account for our experimental paradigm (https://github.com/berenslab/hh_sbi, Table 3).

**Table 3.** Description of 23 extracted electrophysiological features. *To make their distribution more Gaussian, these features are additionally log-transformed, except for the *AP average amp adapt* for which we used the *sigmoid* transform.

| Electrophysiological feature | Description |
| --- | --- |
| *AP threshold* | Membrane voltage at the time where the first derivative of the voltage w.r.t. time reaches a threshold, which elicits the first AP. |
| *AP amplitude* | Height of the 1st AP, measured from threshold to maximum voltage. |
| *AP width* | Width at half height of the 1st AP |
| *AHP* | Afterhyperpolarization. Depth of the membrane voltage drop after the 1st AP, measured from AP threshold. |
| *3rd AP threshold* | Analogous to *AP threshold* but for the 3rd AP. |
| *3rd AP amplitude* | Analogous to *AP amplitude* but for the 3rd AP. |
| *3rd AP width* | Analogous to *3rd AP width* but for the 3rd AP. |
| *3rd AHP* | Analogous to *AHP* but for the 3rd elicited AP. |
| *AP count\** | Number of elicited APs in the current injection window 100 – 700 ms. |
| *AP counts 1st 8th\** | Number of elicited APs in 100 – 175 ms. |
| *AP count 1st quarter\** | Number of APs in 100 – 250 ms. |
| *AP count 1st half\** | Number of APs in 100 – 400 ms. |
| *AP count 2nd half\** | Number of APs in 400 – 700 ms. |
| *AP amp adapt\** | AP amplitude adaptation. 1st elicited *AP amplitude* divided by the amplitude of the 2nd elicited AP. |
| *AP average amp adapt\** | AP average amplitude adaptation. Average ratio of all two consecutive AP heights as calculated by *AP amp adapt* during current injection window. |
| *AP CV\** | Standard deviation divided by the mean of all AP amplitudes of APs elicited during the current injection window. |
| *ISI adapt\** | Interspike interval (ISI) adaptation. ISI: time elapsed between two APs. ISI adapt: ratio of the 2nd ISI (between 2nd and 3rd elicited AP) to the 1st ISI (between 1st and 2nd elicited AP). |
| *ISI CV\** | Standard deviation divided by the mean of all ISIs. |
| *Latency\** | Time it takes to elicit the 1st AP, measured from current stimulation onset to *AP threshold*. |
| *Rest $V_m$ mean* | Mean of the membrane voltage $V_m$ before current stimulation onset 0 – 100 ms. Also called resting membrane potential. |
| *$V_m$ mean* | Mean of the membrane voltage $V_m$ during current stimulation window 100 – 700 ms. |
| *$V_m$ std* | Standard deviation of the membrane voltage $V_m$ during current stimulation window 100 – 700 ms. |
| *$V_m$ skewness* | Skewness of the membrane voltage $V_m$ during current stimulation window 100 – 700 ms. |

### 4.4 Standard neural posterior estimation

Standard NPE applied to HH-based models starts with sampling model parameter sets from a prior distribution *θ p∼* (*θ*), followed by creating synthetic data sets through the simulator or HH-based model, and eventually trains a deep neural density estimator *q*_*ϕ*_ (*θ* |**x**), more specifically in our case, a (masked autoregressive) normalizing flow^57,58^, to learn the probabilistic association between summary statistics **x** derived from simulations and its parameter sets *θ*. Experimental observations **x**_**o**_ can then be fed to the density estimator in order to derive all parameter sets consistent with the data and the prior, i.e. the posterior distribution *q*_*ϕ*_ (*θ* |**x**_*o*_)^25^.

We used the sbi toolbox https://www.mackelab.org/sbi/ to run NPE with different training schedules, including NPE-N, which we explain in the next section.

### 4.5 Dealing with model mismatch in neural posterior estimation

Posterior distributions derived with standard NPE can suffer from model mismatches, that is, when simulations generally fail to adequately cover experimental observations in summary statistic or electrophysiological space. They can become confidently wrong, placing high posterior weight on parameter sets that do not reproduce experimental recordings and low posterior weight to parameter sets that do (Fig. 3b, left). In machine learning jargon, the trained density estimator fails to extrapolate to experimental observations (test data) which are outside of the distribution of the training data.

We experimented with various modifications of NPE in order to make the posterior more robust to the mismatch between model and experimental observations. First, we tried to include only simulations which position themselves close to experimental observations in summary statistic space (Table 1, *Best Euclidean*). Closeness was measured by calculating the Euclidean distance between simulation and observation after standardizing all summary statistics. Second, we introduced different levels of isotropic Gaussian noise to the summaries of those simulations (Table 1, *Add x amplitude noise to ephys*). Third, besides adding noise to summary statistics, we introduced isotropic Gaussian noise to the model parameters with which the close simulations were generated (Table 1, *Add 0*.*05 amplitude noise to ephys and model parameters*). Finally, we experimented with a mixture of non-manipulated close simulations and close simulations with noise added to their summary statistics (Table 1, *Data augmentation*).

Given their performance measures (Table 1), we decided to use NPE with added isotropic Gaussian noise only to the summary statistics of simulations close to experimental observations in summary statistic space. We call the method NPE-N. The noise is of moderate amplitude such that a tradeoff is established between closeness of simulations of MAP estimates to the experimental observations with closeness of simulations from random posterior samples (Table 1, 3rd and 4th column).

In contrast to NPE, NPE-N produced posteriors that give high posterior weight to model parameter sets that both qualitatively and quantitatively produce simulations close to experimental observations.

### 4.6 Feature selection through likelihood marginalization

To analyze which features were informative for constraining the inference procedure, we used Feature Selection through Likelihood Marginalization^37^ (FSLM). To ensure comparable posterior estimates between FSLM and NPE, we trained FSLM with 3 million spiking simulations randomly generated from the prior to which we also introduced isotropic Gaussian noise in their summary statistics. We then drew 1000 samples from the posteriors of each observation *x*_*o*_, for 5 different initializations of FSLM and only selected the 50 experimental observations with smallest average *KL*(*p*_*NPE−N*_(*θ* |*x*_*o*_)|*p*_*FSLM*_(*θ* |*x*_*o*_))^59^ for feature selection. The final ranking was derived from 1000 samples per posterior and is averaged across 10 initializations.

### 4.7 Visualization

We used the *openTSNE*^60^ implementation with default parameters to embed the transcriptomic space with t-SNE to a final two-dimensional representation.

### 4.8 Sparse reduced-rank regression and sparse bottleneck neural networks

To link gene expression data to HH-based model parameters, we used sparse reduced-rank regression (sRRR)^18^. This linear statistical tool reduces the high-dimensional gene space data to a 2-dimensional latent (rank=2), which is maximally predictive of the model parameters. An elastic net penalty was used to select the most relevant genes. As a nonlinear extension to sRRR we also tested the use of sparse bottleneck neural networks (sBNN), that utilize a neural network with bottleneck to predict electrophysiological measurements from transcriptomic space^19^. Analogously to sRRR, a group lasso penalty was used on the weights of the first layer to select most meaningful genes.

## 5 Data availability

Raw electrophysiological recordings are publicly available at https://dandiarchive.org/dandiset/000008/. Further preprocessed data is available either directly in the code repository for this study at https://github.com/berenslab/hh_sbi or on Zenodo (https://doi.org/10.5281/zenodo.7716391). Read counts can be downloaded at https://github.com/berenslab/mini-atlas.

## 6 Code availability

Code to train density neural networks, analyze their performance and produce figures in this manuscript can be found at https://github.com/berenslab/hh_sbi. This code builds upon the simulation-based inference package *sbi*^27^ (https://www.mackelab.org/sbi/), the simulator package *Brian2* (https://brian2.readthedocs.io/en/stable/) and further uses the automatic ephys feature extraction publicly available at https://github.com/berenslab/EphysExtraction, parallel processing package *Pathos*^61^ (https://mmckerns.github.io/project/pathos/wiki.html) and *openTSNE* (https://github.com/pavlin-policar/openTSNE).

## Acknowledgements

We thank Ziwei Huang for discussion. We thank the Deutsche Forschungsgemeinschaft (Heisenberg Professorship BE 5601/8-1 and Excellence Cluster 2064 “Machine Learning — New Perspectives for Science”, ref. 390727645). The work was also funded by the European Union (ERC, “NextMechMod”, ref. 101039115). Views and opinions expressed are however those of the authors only and do not necessarily reflect those of the European Union or the European Research Council Executive Agency. Neither the European Union nor the granting authority can be held responsible for them. Additional support comes from the National Institute of Mental Health and National Institute of Neurological Disorders And Stroke under Award Number U19MH114830. The content is solely the responsibility of the authors and does not necessarily represent the official views of the National Institutes of Health. This work was also supported by the National Institutes of Mental Health grant UM1MH130981 and by the NIH under award number R01 MH109556.

## Author contributions

Conceptualization, P.B., Y.B.; Methodology, Y.B., M.D., P.J.G., J.B., J.H.M., D.K., P.B.; Software, Y.B., M.S.; Formal Analysis, Y.B.; Data Curation, Y.B., F.S.; Writing – Original Draft, Y.B., P.B.; Writing – Review & Editing, all authors; Visualization, Y.B.; Supervision, A.S.T., J.H.M., D.K.,P.B.; Project Administration, P.B.; Funding Acquisition, A.S.T., J.H.M., P.B.

## SUPPLEMENTARY MATERIAL

**Figure S1.**
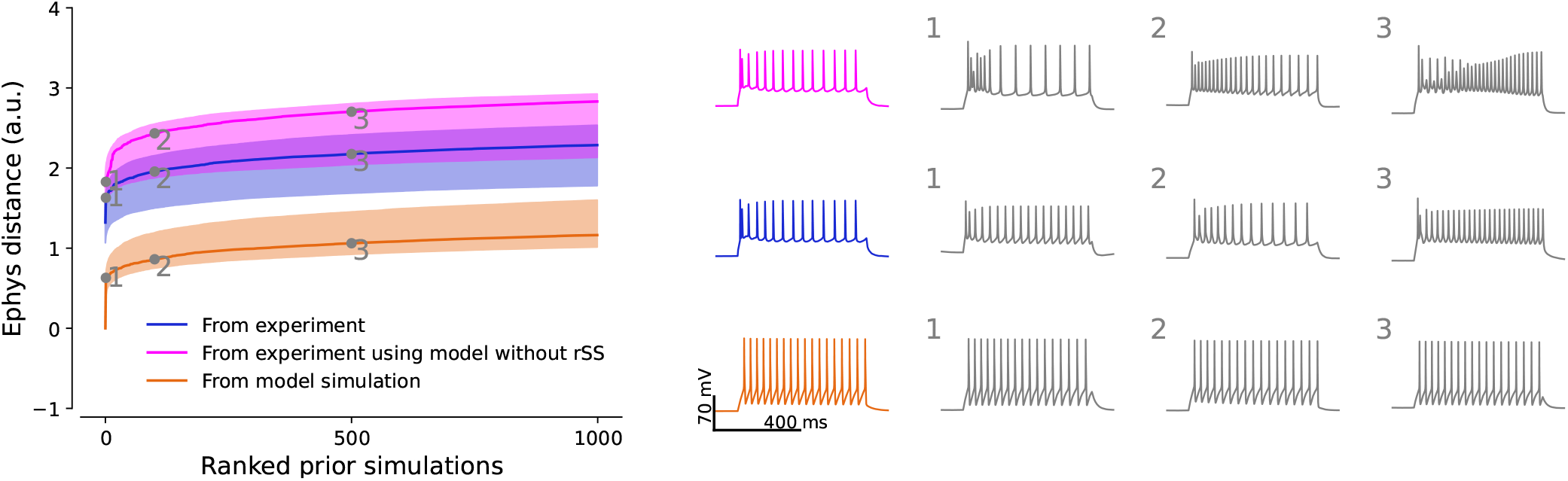
Model misspecification with and without scaling *r*_*SS*_ parameter. Analogous to Fig. 4b, but including model simulations without *r*_*SS*_ parameter.

**Figure S2.**
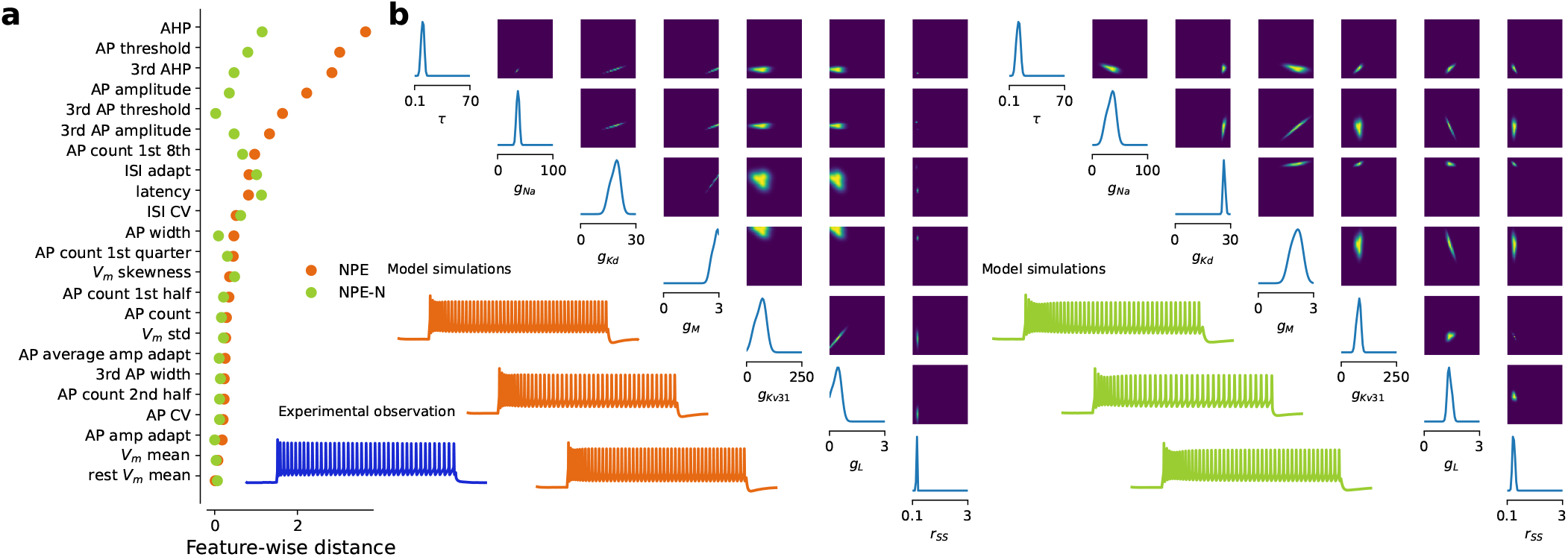
NPE vs NPE-N, illustration 1: fast-spiking *Pvalb Calb1_1* interneuron. Analogous to Fig. 3a,b.

**Figure S3.**
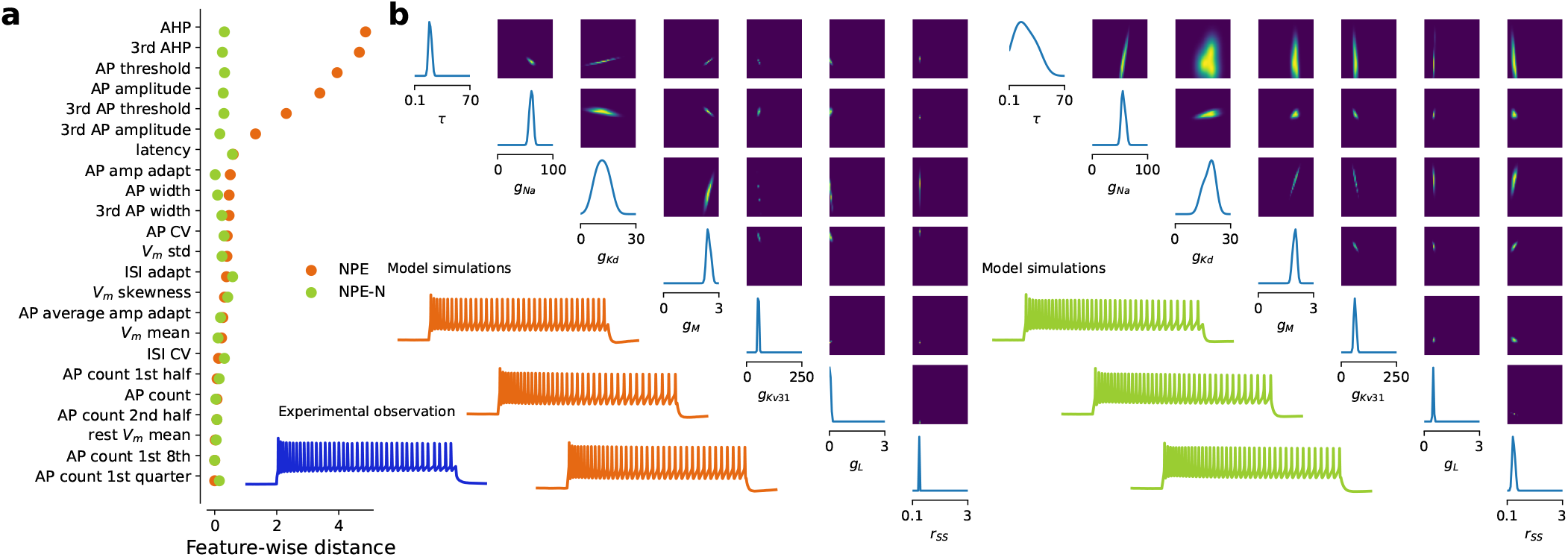
NPE vs NPE-N, illustration 2: *Sst Crhr2_1* interneuron. Analogous to Fig. 3a,b.

**Figure S4.**
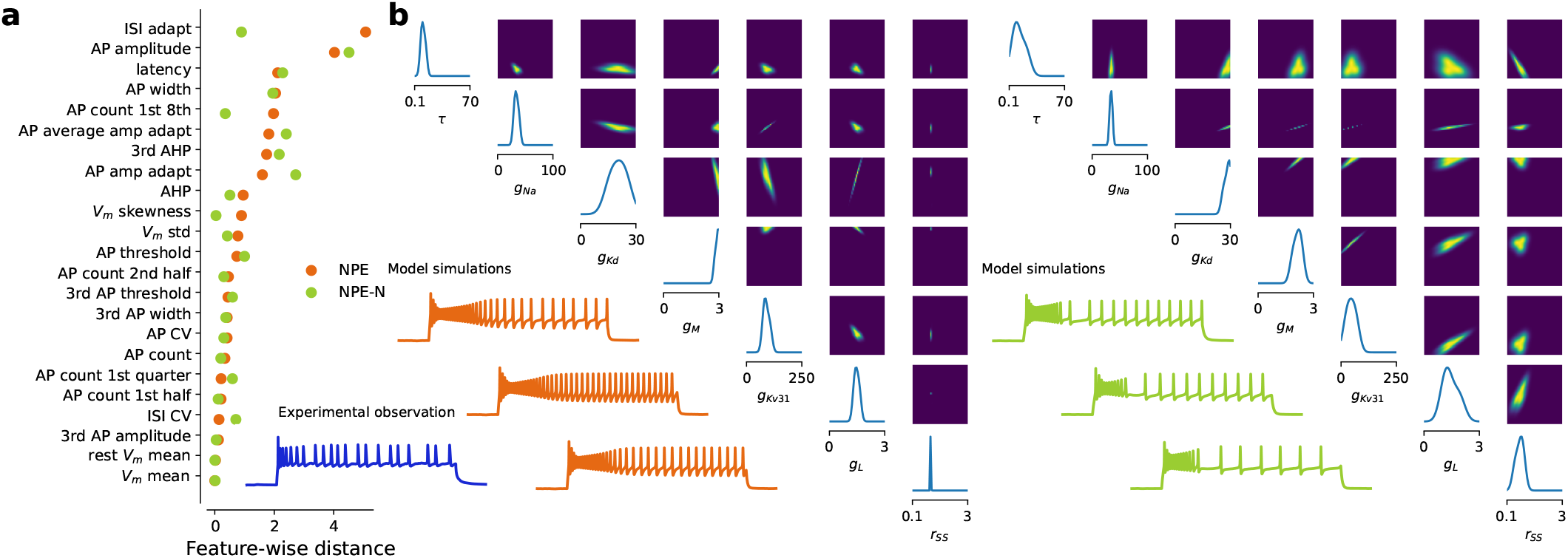
NPE vs NPE-N, illustration 3: *Vip Serpinf1_1* interneuron. Analogous to Fig. 3a,b.

**Figure S5.**
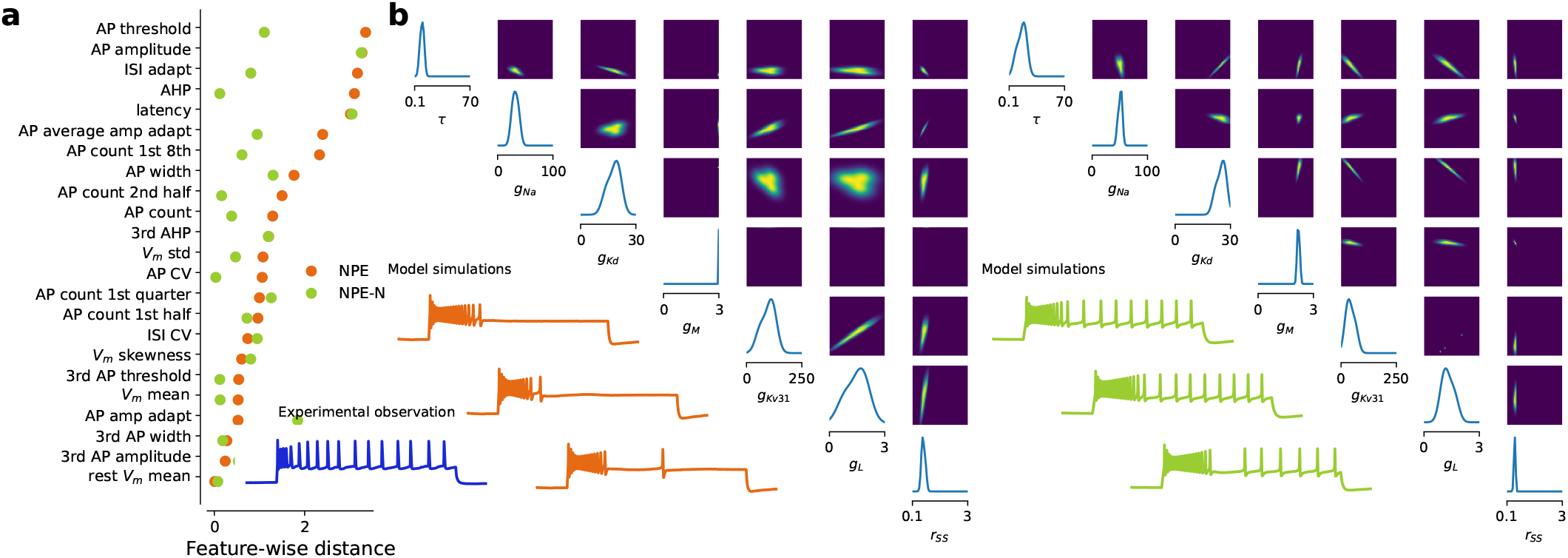
NPE vs NPE-N, illustration 4: *Lamp5 Egln3_1* interneuron. Analogous to Fig. 3a,b.

**Figure S6.**
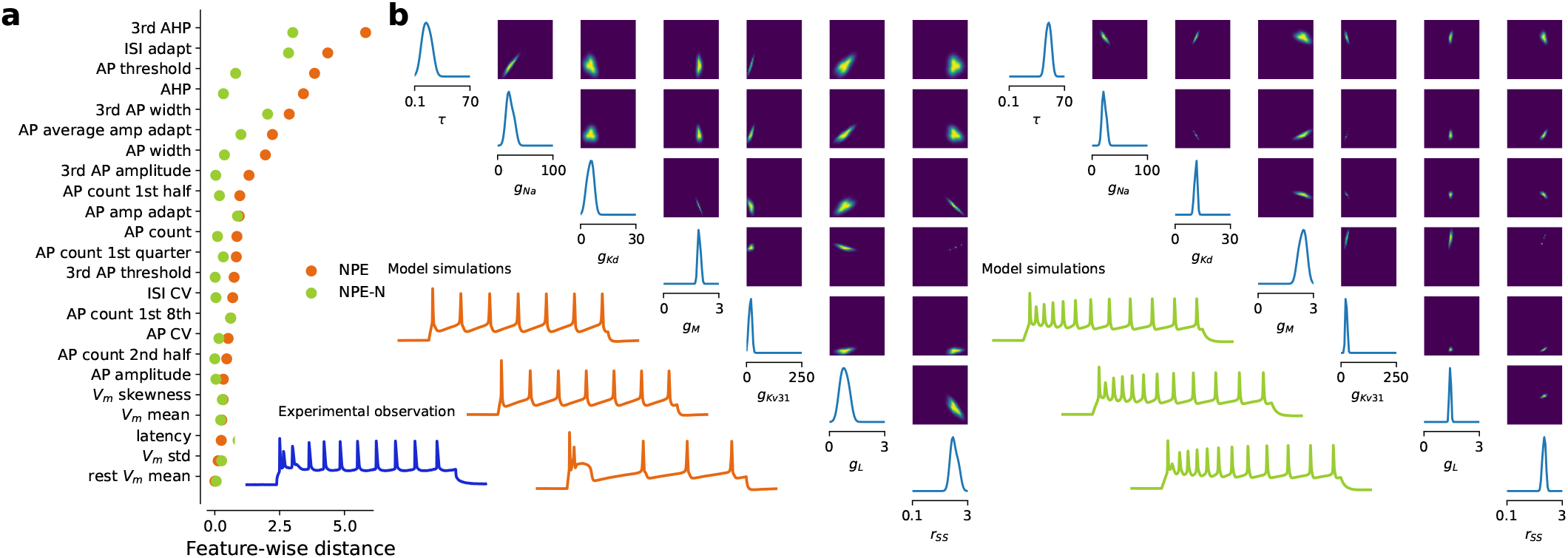
NPE vs NPE-N, illustration 5: *L6 CT Cpa6* pyramidal cell. Analogous to Fig. 3a,b.

**Figure S7.**
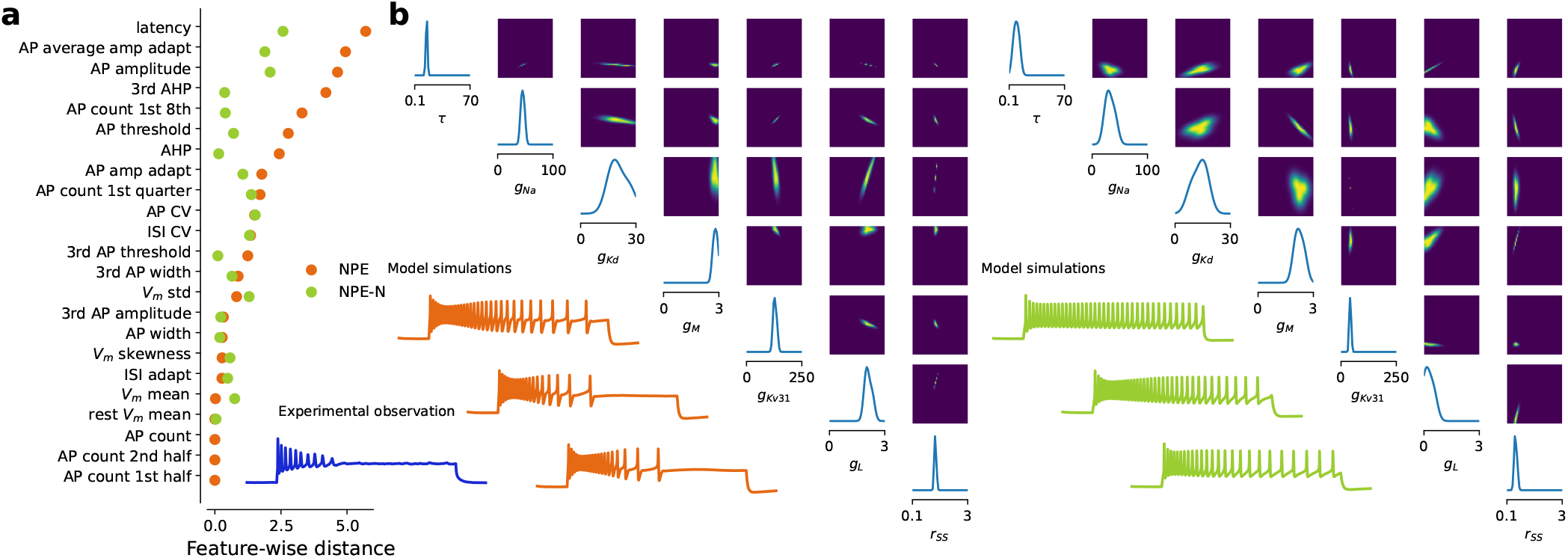
NPE vs NPE-N, illustration 6: *Sst Th_1* interneuron. Analogous to Fig. 3a,b.

**Figure S8.**
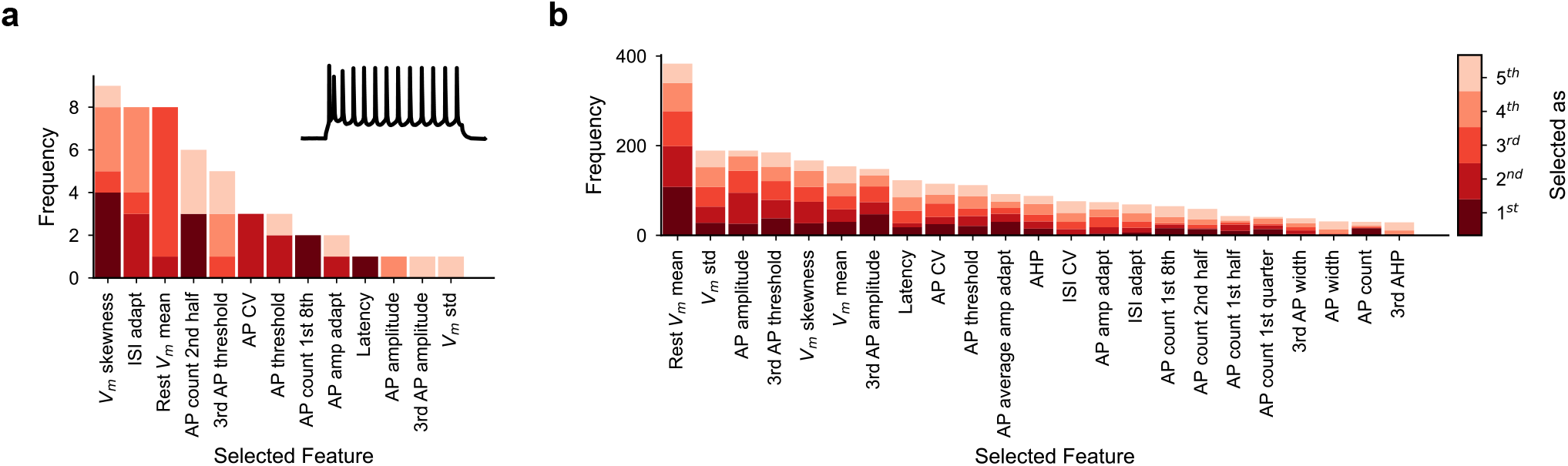
Ranking commonly used electrophysiological features by their ability to constrain posterior estimates. **a** Features are ranked by how often they are strongly constraining the posterior of a Pvalb neuron. Strongly constraining features minimize the KL divergence between posterior estimates subject to all 23 features and estimates considering only five. Important features were selected across 10 repeated runs. Shading indicates the order in which they are selected as part of the top five. Features are ranked in descending order. **b** Summary across all 955 MOp neurons, of which features are strongly constraining the posterior estimates.

## References

1. Zeng, H. & Sanes, J. R. Neuronal cell-type classification: challenges, opportunities and the path forward. Nat. Rev. Neurosci. 18, 530–546 (2017).

2. Douglas, R. J., Martin, K. A. et al. Neuronal circuits of the neocortex. Annu. review neuroscience 27, 419–451 (2004).

3. Harris, K. D. & Shepherd, G. M. The neocortical circuit: themes and variations. Nat. neuroscience 18, 170–181 (2015).

4. Markram, H. et al. Reconstruction and simulation of neocortical microcircuitry. Cell 163, 456–492 (2015).

5. Kepecs, A. & Fishell, G. Interneuron cell types are fit to function. Nature 505, 318–326 (2014).

6. Tremblay, R., Lee, S. & Rudy, B. Gabaergic interneurons in the neocortex: from cellular properties to circuits. Neuron 91, 260–292 (2016).

7. Gouwens, N. W. et al. Classification of electrophysiological and morphological neuron types in the mouse visual cortex. Nat. neuroscience 22, 1182–1195 (2019).

8. Tasic, B. et al. Adult mouse cortical cell taxonomy revealed by single cell transcriptomics. Nat. Neurosci. 19, 335 (2016).

9. Tasic, B. et al. Shared and distinct transcriptomic cell types across neocortical areas. Nature 563, 72 (2018).

10. Zeisel, A. et al. Molecular architecture of the mouse nervous system. Cell 174, 999–1014 (2018).

11. Yao, Z. et al. A transcriptomic and epigenomic cell atlas of the mouse primary motor cortex. Nature 598, 103–110, DOI: 10.1038/s41586-021-03500-8 (2021).

12. Cadwell, C. R. et al. Electrophysiological, transcriptomic and morphologic profiling of single neurons using Patch-seq. Nat. Biotechnol. 34, 199 (2016).

13. Cadwell, C. R. et al. Multimodal profiling of single-cell morphology, electrophysiology, and gene expression using Patch-seq. Nat. Protoc. 12, 2531 (2017).

14. Fuzik, J. et al. Integration of electrophysiological recordings with single-cell rna-seq data identifies neuronal subtypes. Nat. Biotechnol. 34, 175 (2016).

15. Lipovsek, M. et al. Patch-seq: Past, present, and future. J. Neurosci. 41, 937–946 (2021).

16. Gouwens, N. W. et al. Integrated morphoelectric and transcriptomic classification of cortical gabaergic cells. Cell 183, 935–953 (2020).

17. Scala, F. et al. Phenotypic variation of transcriptomic cell types in mouse motor cortex. Nature 598, 144–150 (2021).

18. Kobak, D. et al. Sparse reduced-rank regression for exploratory visualization of paired multivariate data. J. Royal Stat. Soc. Ser. C (2021).

19. Bernaerts, Y., Berens, P. & Kobak, D. Sparse bottleneck neural networks for exploratory non-linear visualization of patch-seq data. ArXiv (2022).

20. Gala, R. et al. A coupled autoencoder approach for multi-modal analysis of cell types. In Advances in Neural Information Processing Systems, 9263–9272 (2019).

21. Gala, R. et al. Consistent cross-modal identification of cortical neurons with coupled autoencoders. Nat. Comput. Sci. 1, 120–127 (2021).

22. Hodkgin, A. L. & Huxley, A. F. A quantitative description of membrane current and its application to conduction and excitation in nerve. The J. Physiol. 117, 500–544 (1952).

23. Pospischil, M. et al. Minimal hodgkin-huxley type models for different classes of cortical and thalamic neurons. Biol. Cybern. 99, 427–441, DOI: 10.1007/s00422-008-0263-8 (2008).

24. Hay, E., Hill, S., Schürmann, F., Markram, F. & Segev, I. Models of neocortical layer 5b pyramidal cells capturing a wide range of dendritic and perisomatic active properties. PLOS Comput. Biol. 7, DOI: 10.1371/journal.pcbi.1002107 (2011).

25. Gonçalves, P. J. et al. Training deep neural density estimators to identify mechanistic models of neural dynamics. eLife 9, DOI: 10.7554/eLife.56261 (2020).

26. Greenberg, D., Nonnenmacher, M. & Macke, J. Automatic posterior transformation for likelihood-free inference. In Chaudhuri, K. & Salakhutdinov, R. (eds.) Proceedings of the 36th International Conference on Machine Learning, vol. 97 of Proceedings of Machine Learning Research, 2404–2414 (PMLR, 2019).

27. Tejero-Cantero, A. et al. sbi: A toolkit for simulation-based inference. J. Open Source Softw. 5, 2505, DOI: 10.21105/joss.02505 (2020).

28. Gouwens, N. W. et al. Systematic generation of biophysically detailed models for diverse cortical neuron types. Nat. Commun. 9, DOI: 10.1038/s41467-017-02718-3 (2018).

29. Nandi, A. et al. Single-neuron models linking electrophysiology, morphology, and transcriptomics across cortical cell types. Cell Reports 40, 111176, DOI: 10.1016/j.celrep.2022.111176 (2022).

30. Ward, D., Cannon, P., Beaumont, M., Fasiolo, M. & Schmon, S. M. Robust neural posterior estimation and statistical model criticism. In Oh, A. H., Agarwal, A., Belgrave, D. & Cho, K. (eds.) Advances in Neural Information Processing Systems (2022).

31. Cannon, P., Ward, D. & Schmon, S. M. Investigating the impact of model misspecification in neural simulation-based inference, DOI: 10.48550/ARXIV.2209.01845 (2022).

32. Schmitt, M., Bürkner, P.-C., Köthe, U. & Radev, S. T. Detecting model misspecification in amortized bayesian inference with neural networks. In Köthe, U. & Rother, C. (eds.) Pattern Recognition, 541–557 (Springer Nature Switzerland, Cham, 2024).

33. David T. Frazier, C.D., David J. Nott & Kohn, R. Bayesian inference using synthetic likelihood: Asymptotics and adjustments. J. Am. Stat. Assoc. 118, 2821–2832, DOI: 10.1080/01621459.2022.2086132 (2023). https://doi.org/10.1080/01621459.2022.2086132.

34. Stimberg, M., Brette, R. & Goodman, D. F. Brian 2, an intuitive and efficient neural simulator. eLife 8, DOI: 10.7554/eLife.47314 (2019).

35. Papamakarios, G. & Murray, I. Fast ε-free inference of simulation models with bayesian conditional density estimation. In Lee, D., Sugiyama, M., Luxburg, U., Guyon, I. & Garnett, R. (eds.) Advances in Neural Information Processing Systems, vol. 29 (Curran Associates, Inc., 2016).

36. Cranmer, K., Brehmer, J. & Louppe, G. The frontier of simulation-based inference. PNAS 117, 30055–30062, DOI: 10.1073/pnas.1912789117 (2020).

37. Beck, J., Deistler, M., Bernaerts, Y., Macke, J. & Berens, P. Efficient identification of informative features in simulation-based inference. In Advances in Neural Information Processing Systems (Curran Associates, Inc., 2022).

38. Apicella, A.j. & Marchionni, I. Vip-expressing gabaergic neurons: Disinhibitory vs. inhibitory motif and its role in communication across neocortical areas. Front. Cell. Neurosci. 16, DOI: 10.3389/fncel.2022.811484 (2022).

39. Traub, R. D. & Miles, R. Neuronal networks of the Hippocampus (Cambridge University Press, 1991).

40. Prinz, A. A., Bucher, D. & Marder, E. Similar network activity from disparate circuit parameters. Nat. neuroscience 7, 1345–1352 (2004).

41. Deistler, M., Macke, J. H. & Gonçalves, P. J. Energy-efficient network activity from disparate circuit parameters. Proc. Natl. Acad. Sci. 119, e2207632119 (2022).

42. Frazier, D. T. & Drovandi, C. Robust approximate bayesian inference with synthetic likelihood. J. Comput. Graph. Stat. 30, 958–976, DOI: 10.1080/10618600.2021.1875839 (2021). https://doi.org/10.1080/10618600.2021.1875839.

43. Lause, J., Berens, P. & Kobak, D. Analytic pearson residuals for normalization of single-cell rna-seq umi data. Genome Biol. 22, 258, DOI: 10.1186/s13059-021-02451-7 (2021).

44. Schölkopf, B. Causality for machine learning. In Probabilistic and Causal Inference: The Works of Judea Pearl, 765–804 (Association for Computing Machinery, 2022).

45. Buccitelli, C. & Selbach, M. mrnas, proteins and the emerging principles of gene expression control. Nat. Rev. Genet. 21, 630–644 (2020).

46. Srivastava, H. et al. Protein prediction models support widespread post-transcriptional regulation of protein abundance by interacting partners. PLOS Comput. Biol. 18, e1010702 (2022).

47. Schulz, D. J.Goaillard, J.-M. & Marder, E. E. Quantitative expression profiling of identified neurons reveals cell-specific constraints on highly variable levels of gene expression. Proc Natl Acad Sci U S A 104, 13187–13191 (2007).

48. Zur, H. & Tuller, T. Predictive biophysical modeling and understanding of the dynamics of mrna translation and its evolution. Nucleic acids research 44, 9031–9049 (2016).

49. Kotaleski, J. H. & Blackwell, K. T. Modelling the molecular mechanisms of synaptic plasticity using systems biology approaches. Nat. Rev. Neurosci. 11, 239–251 (2010).

50. Holt, C. E., Martin, K. C. & Schuman, E. M. Local translation in neurons: visualization and function. Nat. structural & molecular biology 26, 557–566 (2019).

51. Morisaki, T. & Stasevich, T. J. Quantifying single mrna translation kinetics in living cells. Cold Spring Harb. perspectives biology 10, a032078 (2018).

52. Tutucci, E., Livingston, N. M., Singer, R. H. & Wu, B. Imaging mrna in vivo, from birth to death. Annu. review biophysics 47, 85–106 (2018).

53. Colbert, C. M. & Pan, E. Ion channel properties underlying axonal action potential initiation in pyramidal neurons. Nat. Neurosci. 5, 533–538, DOI: 10.1038/nn0602-857 (2002).

54. Yamada, W. M., Koch, C. & Adams, P. R. Methods in neuronal modeling: From synapses to networks (MIT press, 1989).

55. Rettig, J. et al. Characterization of a shaw-related potassium channel family in rat brain. The EMBO journal 11, 2473–2486, DOI: j.1460-2075.1992.tb05312.x (1992).

56. Reuveni, I., Friedman, A., Amitai, Y. & Gutnick, M. Stepwise repolarization from ca2+ plateaus in neocortical pyramidal cells: evidence for nonhomogeneous distribution of hva ca2+ channels in dendrites. J. Neurosci. 11, 4609–4621, DOI: 10.1523/JNEUROSCI (1993).

57. Papamakarios, G., Nalisnick, E., Rezende, D. J., Mohamed, S. & Lakshminarayanan, B. Normalizing flows for probabilistic modeling and inference. J. Mach. Learn. Res. 22, 1–64 (2021).

58. Papamakarios, G., Pavlakou, T. & Murray, I. Masked autoregressive flow for density estimation. In Advances in Neural Information Processing Systems, 2335–2344 (2017).

59. Jiang, B. Approximate bayesian computation with kullback-leibler divergence as data discrepancy. In Storkey, A. J. & Pérez-Cruz, F. (eds.) International conference on artificial intelligence and statistics, AISTATS 2018, vol. 84 of Proceedings of machine learning research, 1711–1721 (PMLR, 2018).

60. Poličar, P. G., Stražar, M. & Zupan, B. opentsne: A modular python library for t-sne dimensionality reduction and embedding. J. Stat. Softw. 109, 1–30, DOI: 10.18637/jss.v109.i03 (2024).

61. Michael M. McKerns, Leif Strand, Tim Sullivan, Alta Fang & Michael A.G. Aivazis. Building a Framework for Predictive Science. In Stéfan van der Walt & Jarrod Millman (eds.) Proceedings of the 10th Python in Science Conference, 76 – 86, DOI: 10.25080/Majora-ebaa42b7-00d (2011).

